# Dual lineages of Langerhans cells cooperate to restore the immune barrier after skin injury

**DOI:** 10.64898/2026.05.25.727646

**Authors:** Axel D. Schmitter-Sánchez, Nicholas Basista, Sudhanshu Mishra, Hyeri Kim, Audrey Bench, Catherine Matte-Martone, Min-Soo Seo, Gun Woo Lee, Sangbum Park

## Abstract

Langerhans cells (LCs) are key immune sentinels of the epidermis. How this network reorganizes to safeguard epidermal immunity after injury has remained unclear. Here, we uncover a previously unrecognized two-lineage program of LC repopulation during wound repair. Classically, tissue-resident embryonically derived LCs (eLCs) migrate to lymph nodes in response to antigens. In contrast, we find that injury triggers nearby eLCs to migrate into wounds, providing immediate coverage. In parallel, circulating monocytes infiltrate the skin and differentiate into long-lived monocyte-derived LCs (mLCs) that integrate stably into the network. We identify the chemokine receptor CXCR2 as a novel regulator of eLC migration into wounds, distinct from the CXCR4/CCR7 pathways mediating LC egress to lymph nodes. Pharmacological inhibition of CXCR2 impairs directional eLC migration and is accompanied by increased mLC infiltration, preserving immune barrier density. These findings reveal a coordinated and flexible two-lineage repair program that ensures robust restoration of epidermal immunity.

## Main

The skin epidermis, the outermost layer of the body, functions as the primary barrier against the external environment. Together, a multi-layer structure of epithelial cells and a network of tissue-resident immune cells provide physical and immunological protection, respectively^1,2^. When the skin is damaged, epithelial cells around the wound promptly coordinate migration and proliferation to restore the barrier in a critical process called re-epithelialization, which is followed by a prolonged remodeling phase that returns tissue homeostasis^3,4^. Tissue-resident immune cells also reestablish the immunological barrier after injury; however, the underlying mechanism remains unclear.

Langerhans cells (LCs) represent the predominant immune cell population in the epidermis. LCs share a common ontogeny with dendritic cells and macrophages, continuously surveying the epidermal microenvironment^5^. Upon encountering antigens, LCs depart the epidermis to act as antigen-presenting cells (APCs) by migrating to nearby lymph nodes to prime adaptive immunity^6^. Beyond this canonical dendritic cell role, LCs also display macrophage-like features, contributing to skin innervation, angiogenesis, and repair, reflecting a dual identity shaped by both intrinsic and niche-derived signals ^7^.

Given their vital role in skin immunity, understanding how LCs develop and maintain their presence in the epidermis has been the focus of extensive research. Under homeostatic conditions, embryonically derived LCs (eLC) are maintained by local self-renewal^8–10^. When LC density is severely reduced, progenitor cells are recruited into the epidermis and differentiate into LCs to repopulate the lost network^11–14^. Ultraviolet-induced depletion of LCs leads to an initial recruitment of Gr1^hi^ monocytes that give rise to “short-term” LCs, later replaced by unidentified precursor cells that generate “long-term” LCs^13^. In contrast, LC elimination by immune-mediated pathology triggers recruitment of monocytes to the epidermis, which directly give rise to long-lived monocyte-derived LCs (mLC)^12^. Hair follicle epithelial cells play a critical role in recruiting the monocytes, acting as an instructive niche where Jagged–Notch signaling drives early commitment to the LC lineage^7^.

Despite these insights, how the LC network is rebuilt after mechanical disruption of the epidermis, such as in wound healing, remains poorly understood. A few previous studies have reported LC repopulation within the newly formed epithelium after injury^15–17^, implicating circulating progenitors, but the mechanism was not clear. To address this gap, we combined intravital imaging and lineage tracing to track the fate of LCs during wound repair.

We discovered that most tissue-resident eLCs near the wound do not exit the epidermis but instead migrate toward the wound during re-epithelialization. Following migration, these LCs exhibit enhanced proliferation. Mechanistically, our data suggest that signaling through chemokine receptor CXCR2 promotes LC migration towards the wound. At the same time, monocytes infiltrate the wounded epidermis, differentiate into mLCs, and become stably integrated into the LC network. Transcriptomic characterization of these two LC populations revealed distinct gene signatures: newly differentiated mLCs are enriched in genes associated with immune tolerance, whereas pre-existing eLCs are enriched in pathways related to tissue regeneration. These data suggest their potentially divergent roles in wound repair. Overall, our findings reveal that the reconstruction of the LC immunological barrier after tissue damage entails the cooperation of two different lineages of LCs.

## Results

### LCs from the nearby epidermis migrate into the wound

To investigate the longitudinal changes in LCs after injury in live mice, we used our established intravital imaging approach using fluorescent reporter mice (Figure S1A, B)^3,18–20^. The mouse line we generated carries the *Lang-EGFP* allele^21^, which labels the cytoplasm of LCs with EGFP, and the *K14-H2B-mCherry* allele^22^, which labels epithelial nuclei with mCherry. We created a full-thickness wound on the dorsal skin of the mouse ear using a 1-mm punch biopsy tool, as described previously^3^. By five days post-injury, epithelial cells had fully covered the wound area through re-epithelialization. Notably, we also observed LCs within the re-epithelialized area, consistent with previous reports (Figures 1A)^15–17^.

**Figure 1.**
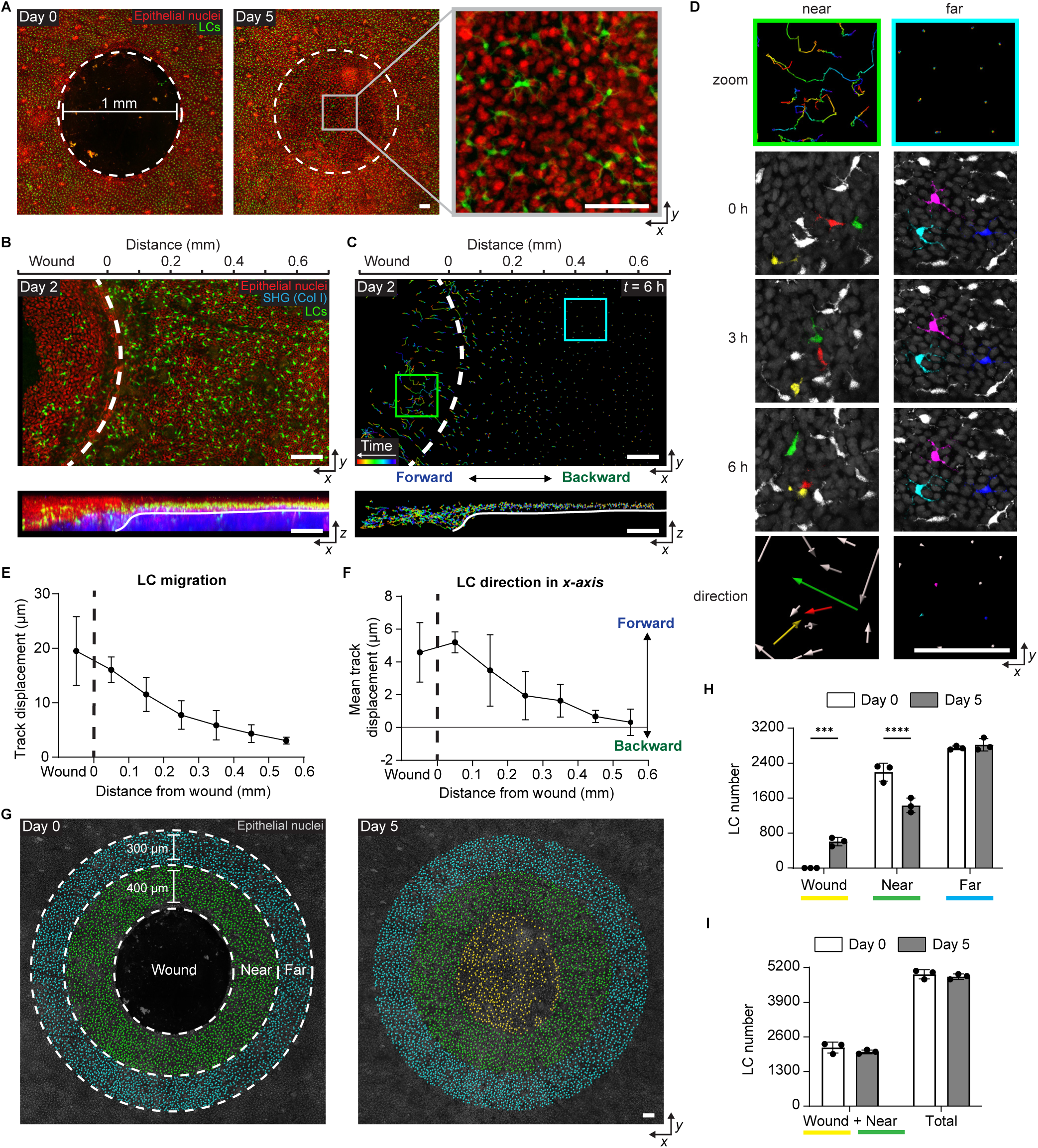
Langerhans cells migrate towards the wound during re-epithelization. **A**, Revisit multi-photon *in-vivo* microscopy images of a 1 mm wound from the same mouse. Image shows *x-y* view of epithelial cells (red nuclei) and LCs (green) in the epidermis. Dashed line indicates initial wound boundary. *Left*: Day of wound induction (Day 0). *Middle*: 5 days after wound induction. *Right*: zoomed view of the wound center at Day 5. Representative images from 3 mice. Scale bars, 100 µm. **B**, Time-lapse image of epithelial cells (red nuclei) and LCs (green) 2 days after wound induction. Dashed line, initial wound boundary. Solid line, basal membrane separating epidermis from dermis. *Top*: *x-y* view. *Bottom*: *x-z* view shows the epidermis (red) and dermis (collagen SHG, blue). Representative images from 3 mice. Scale bars, 100 µm. **C**, Imaris track analysis of LCs (**B**) 2 days after wound induction. Colors project time (blue, 0h; red, 6h). *Top*: *x-y* view. *Bottom*: *x-z* view. Representative images from 3 mice. Scale bars, 100 µm. **D**, *Top*: zoomed migration tracks from (**C**). The green frame is from the wound leading-edge epithelial migration zone, and the teal frame is from the epithelial proliferation zone^3^. *Middle*: time-lapse frames show the movement of individually colored LCs across 6 hours. Epithelial cell nuclei in gray. Other LCs in white. *Bottom*: vector arrows show the general movement direction of the matching color LC. Representative images from 3 mice. Scale bars, 100 µm. **E**, Mean total displacement of individual LC tracks over 6h plotted as a function of distance from the wound. *n* = 3 mice. **F**, Mean track displacement in the *x* axis of individual LC tracks over 6h plotted as a function of distance from the wound. Calculated by comparing the start and end values in the *x* axis of each track. Positive change indicates movement towards the wound. *n* = 3 mice. **E-F**, Imaging performed 2 days after wound induction. Dashed line, initial wound boundary. The displacements of migrating cell tracks were averaged every 100 µm from the initial wound. Data are mean ± s.d. **G**, Imaris cell count analysis of LCs (spots) at Day 0 (*left*) and 5 days after wound induction (*right*). Dashed lines separate LCs into 3 zones: wound (yellow spots), near (0-400 µm from the wound edge, green spots), and far (400-700 µm from the wound edge, teal spots). Epithelial cell nuclei are shown in gray. Representative images from 3 mice. Scale bars, 100 µm. **H**, Mean LC number comparing cell density between Day 0 and 5 days after wound induction according to the 3 zones established in (**G**). *n* = 3 mice. **I**, Mean LC number comparing the cell density change from the addition of the wound and near zones between Day 0 and 5 days after wound induction (*left bars*). Total change in LC density across all 3 zones between Day 0 and 5 days after wound induction (*right bars*). *n* = 3 mice. **H-I**, Data analyzed using paired two-way ANOVA; data are mean ± s.d. with each dot representing individual mice. *** *P* < 0.001, **** *P* < 0.0001.

Circulating monocytes have been reported to transiently differentiate into epidermal LCs in response to depletion^11,13^. This mechanism has been proposed to contribute to LC reconstitution after injury, but the rapid appearance of LCs in our wound model (Day 5) occurs too quickly to be fully explained by monocyte infiltration and differentiation^12,13^. Therefore, we hypothesized that LCs adjacent to the wound edge migrate toward the wound site. To test this hypothesis, we performed time-lapse imaging at two days post-injury. Interestingly, LCs near the wound exhibited dynamic movement, whereas those farther from the wound remained stationary (Figure 1B-D and Video S1). This spatial organization of LC behavior parallels our previous findings that epithelial cells establish distinct territories during re-epithelialization^3^, whereby epithelial cells near the wound collectively migrate during re-epithelialization (Figure S1C-E).

However, migration track analysis revealed that, unlike epithelial cells, LC movement was irregular and independent of neighboring LCs and epithelial cells (Figure 1C, D). To determine if LCs actively migrate toward the wound or merely patrol the local environment, we conducted automated quantitative tracking and directional analysis. Despite individual variations in movement direction and displacement, the majority of LCs near the wound exhibited directed migration toward the wound (Figure 1E, F). This result suggests that LCs near the wound edge actively migrate to the re-epithelialized area immediately after injury.

Given that the primary function of LCs is to deliver antigen to naïve T cells in lymph nodes, we initially expected that many LCs near the wound would exit the epidermis. However, time-lapse imaging revealed that almost all LCs near the wound remained within the epidermis. To quantify changes in LC density after injury, we measured LC numbers at different distances from the wound immediately after injury (Day 0) and after wound closure (Day 5) (Figure 1G). LC density after re-epithelialization was significantly reduced near the wound but was unchanged farther from the wound, where LCs had remained stationary after injury (Figure 1H). Because our time-lapse data showed LCs near the wound move into the wound site during re-epithelialization, we reasoned that this local drop in LC density reflected redistribution. Indeed, an area including both “wound” and “near” regions at Day 5 contained more than 92.8% of LCs present in the “near” region at Day 0 (Figure 1I). These data suggest that LCs near the wound display dynamic mobility immediately after injury to repopulate the wound site during re-epithelialization.

These findings reveal an unanticipated migratory behavior of LCs during wound healing, distinct from their well-known lymph node trafficking. We propose that local LCs safeguard epidermal immunity by migrating into wounds to restore the barrier.

## Epithelial migration is dispensable for LC mobility near the wound

Our previous study demonstrated a gradient of decreasing epithelial cell displacement as the distance from the wound edge increases^3^. Intriguingly, LC migration exhibited a similar spatial pattern to that of epithelial cells (Figures 1E and S1E). LCs are tightly connected to adjacent epithelial cells via cell adhesion molecules such as E-cadherin and EpCAM^23,24^. Given the parallel spatial distribution and physical adhesion between epithelial cells and LCs, we considered that epithelial migration influences LC migration.

To test this, we specifically inhibited epithelial cell mobility in vivo via conditional epithelial loss-of-function of the Rho GTPase Rac1^3^. We generated an epithelial-specific Rac1 conditional knockout in our fluorescent reporter mouse line, using a keratin 14 promoter-driven inducible CreER system activated by tamoxifen injection (*K14-CreER; Rac1^−/−^*, hereafter referred to as Epi-Rac1 KO; Figure S2A). We have previously shown that this model effectively impairs epithelial cell migration during wound repair (Figure S2B, C)^3^. Consistent with this established phenotype, time-lapse imaging two days post-injury revealed markedly reduced epithelial cell movement in Epi-Rac1 KO mice relative to littermate controls (*K14-CreER; Rac1^+/−^*; Figure 2A, C).

**Figure 2.**
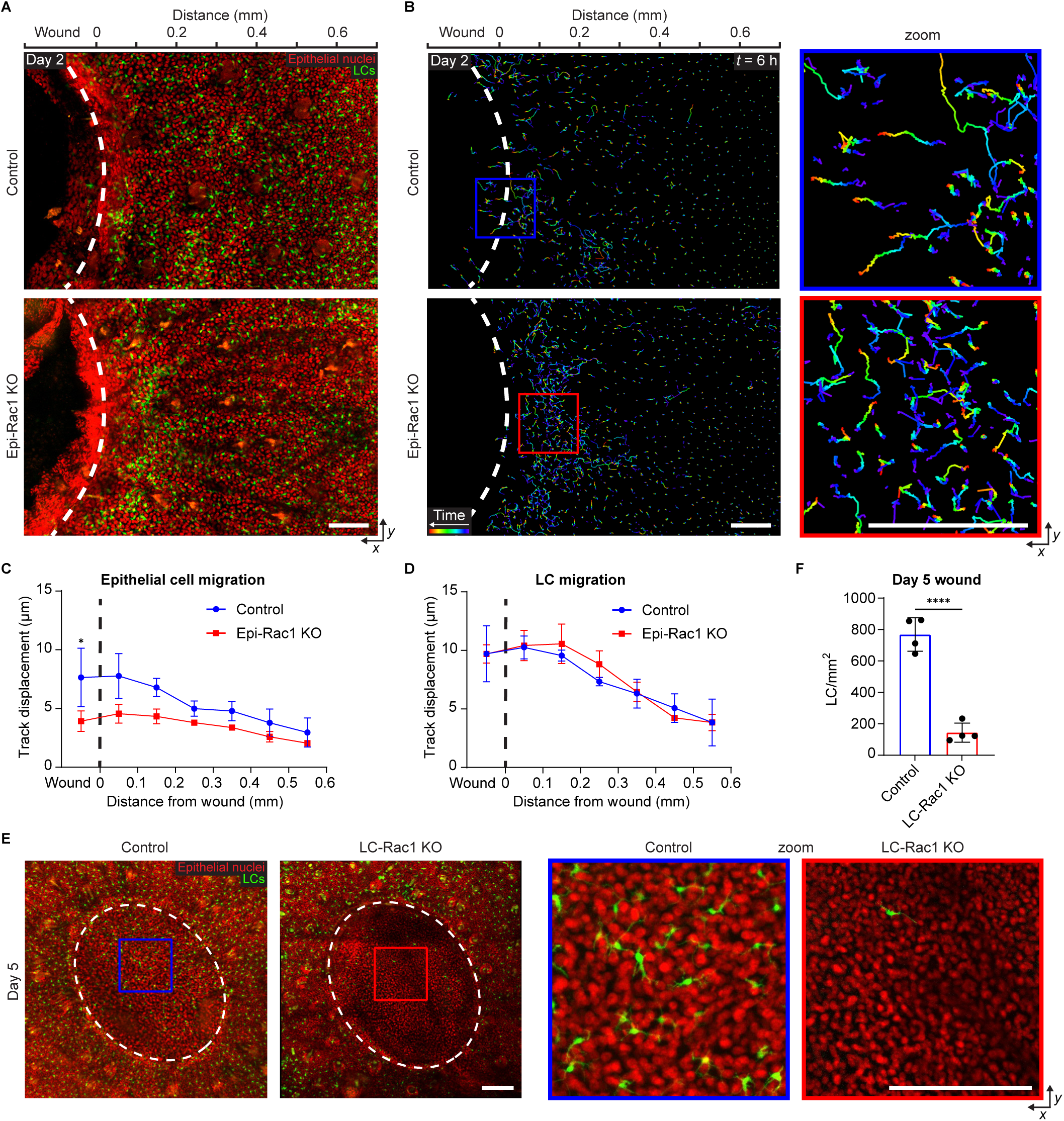
Langerhans cell mobility is independent of epithelial cell migration. **A**, Time-lapse *x-y* view of epithelial cells (red nuclei) and LCs (green) 2 days after wound induction. *Top*: control mouse. *Bottom*: Epi-*Rac1 KO* mouse. Dashed line indicates initial wound boundary. Representative images from 3 mice per group. Scale bars, 100 µm. **B**, Imaris *x-y* view track analysis of LCs (**A**) 2 days after wound induction. Colors project time (blue, 0h; red, 6h). *Top*: control mouse. *Bottom*: Epi-*Rac1 KO* mouse. *Right*: zoomed migration tracks from near the wound edge. Dashed line indicates initial wound boundary. Representative images from 3 mice per group. Scale bars, 100 µm. **C**, Mean total displacement of individual epithelial cell tracks from control and Epi-*Rac1 KO* mice over 6h plotted as a function of distance from the wound. *n* = 3 mice per group **D**, Mean total displacement of individual LC tracks from control and Epi-*Rac1 KO* mice over 6h plotted as a function of distance from the wound. *n* = 3 mice per group **C-D**, Imaging performed 2 days after wound induction. Dashed line, initial wound boundary. The displacements of migrating cell tracks were averaged every 100 µm from the initial wound. Data analyzed using unpaired two-way ANOVA; data are mean ± s.d. * *P* < 0.05. **E**, *In-vivo* microscopy images shows *x-y* view of epithelial cells (red nuclei) and LCs (green) in the epidermis 5 days after wound induction. Dashed line indicates initial wound boundary. *Left*: control mouse and LC-*Rac1 KO* mouse. *Right*: zoomed view of the wound center from 5 days after wound induction. Representative images from 4 mice per group. Scale bars, 200 µm. **F**, Mean LC number inside the wound epidermis from control and LC-*Rac1 KO* mice. Imaging was performed 5 days after wound induction. LC density normalized to the individual mouse wound area quantified. Data analyzed using unpaired two-tailed *t*-test; *n =*4 mice per group; data are mean ± s.d. with each dot representing individual mice. **** *P* < 0.0001.

Notably, LCs near the wound in Epi-Rac1 KO mice still exhibited dynamic movement, despite the lack of epithelial cell mobility (Figure 2B and Video S2). Quantification of LC migration revealed that the overall displacement of LCs did not significantly differ between Epi-Rac1 KO and control mice (Figure 2D).

These results suggest that LC mobility is triggered by the wound and does not require the movement of neighboring epithelial cells.

To determine whether LC movement is cell-autonomous, we established a conditional knockout of Rac1 in LCs using a Langerin promoter-driven inducible CreER, as previously described (*huLangerin-CreER; Rac1^−/−^*, hereafter referred to as LC-Rac1 KO; Figure S2A)^20^. In littermate controls (*huLangerin-CreER; Rac1^+/+^*), LCs migrated into the re-epithelialized epidermis by Day 5 post-injury, whereas in LC-Rac1 KO mice, LCs barely entered the re-epithelialized region Figure 2E, F).

Together, these findings demonstrate that LC mobility is stimulated by wounding, cell-autonomous, and dependent on Rac1 within LCs. Although epithelial cells and LCs migrate in adjacent wound areas, epithelial migration is not required for LC motility per se.

## Temporally coordinated proliferation and loss restore Langerhans cell homeostasis after injury

Our data so far show that adjacent LCs contribute to restoring the immunological barrier by migrating toward the wound. The movement of epidermal-resident LCs facilitated the rapid redistribution of LCs to cover the injured area. However, since LC redistribution occurred without significant proliferation or influx of new cells during re-epithelialization, LC density in and near the wound was initially lower than normal (Figure 1H). To assess full recovery, we repeatedly imaged the same wounds over time to track when LC density returned to baseline (Figure 3A).

**Figure 3.**
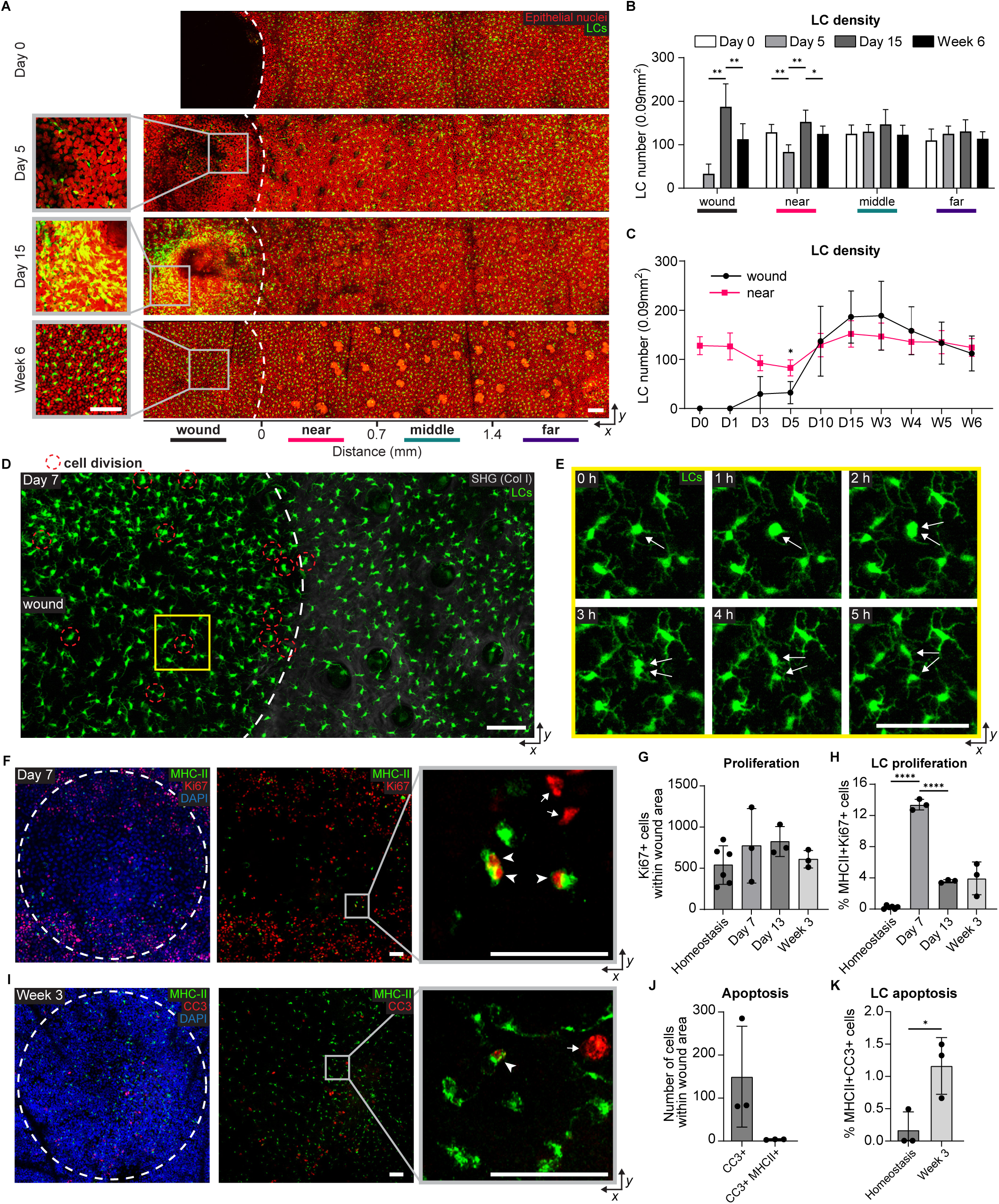
Langerhans cell density changes during the remodeling phase. **A**, Revisit multi-photon *in-vivo* microscopy images of a 1 mm wound from the same mouse. Images show *x-y* view of epithelial cells (red nuclei) and LCs (green) in the epidermis at 0, 5, 15 days, and 6 weeks after wound induction. Dashed line indicates initial wound boundary. *Left*: zoomed view of the wound center. *Right*: revisit images showing the classification of several zones according to their distance from the wound: wound, near, middle, and far. Representative images from 6 mice. Scale bars, 100 µm. **B**, Mean LC number comparing cell density changes within designated zones established in (**A**) at 0, 5, 15 days, and 6 weeks after wound induction. *n* = 6 mice. **C**, Timeline of LC number comparing cell density changes between the wound and near zones. *n* = 6 mice. **B-C**, Data analyzed using unpaired two-way ANOVA; data are mean ± s.d. * *P* < 0.05, ** *P* < 0.01. **D**, Time-lapse image in *x-y* view of proliferative LCs (green) at 7 days after wound induction. Dermis SHG collagen is shown in gray. Red dashed circles indicate diving LCs. **E**, Time-lapse frames from yellow highlighted area in (**D**) show LC division across 5 hours. White arrows indicate actively diving LC. **D-E**, Representative images from 3 mice. White dashed circle/line indicates initial wound boundary. Scale bars, 50 µm. **F**, Confocal immunofluorescent images of cell proliferation at the wound epidermis 7 days after wound induction. Images show *x-y* view of LCs (green, MHC-II), proliferation (red, Ki67), and cell nuclei (blue, DAPI). *Left*: composite image. *Middle*: MHC-II and Ki67 positive cells. *Right*: zoomed example of proliferative LC. MHC-II, major histocompatibility complex class II. White arrows show proliferative cells. White arrowheads show proliferative LCs. Representative images from 4 mice. The white dashed line indicates the initial wound boundary. Scale bars, 50 µm. **G**, Timeline of mean proliferative (Ki67+) cell density at the wound during healing (*n* = 3 mice) and in homeostasis (*n* = 6 mice). **H**, Timeline of percentage of proliferative LCs (MHC-II+Ki67+) at the wound during healing (*n* = 3 mice) and in homeostasis (*n* = 6 mice). **I**, Confocal immunofluorescent images of cell apoptosis at the wound epidermis 3 weeks after wound induction. Images show *x-y* view of LCs (green, MHC-II), apoptosis (red, CC3), and cell nuclei (blue, DAPI). *Left*: composite image. *Middle*: MHC-II and CC3 positive cells. *Right*: zoomed example of apoptotic LC. CC3, cleaved caspase-3. The white arrow shows an apoptotic cell. White arrowhead shows an apoptotic LC. Representative images from 4 mice. The white dashed line indicates the initial wound boundary. Scale bars, 50 µm. **J**, Mean number of apoptotic cells (CC3+) and apoptotic LCs (CC3+MHC-II+) at the wound 3 weeks after wound induction. *n* = 3 mice. **K**, Percentage of apoptotic LCs (CC3+MHC-II+) during homeostasis and at 3 weeks after wound induction. *n* = 3 mice. **J-K**, Wound area quantified 0.49 mm^2^ per mouse. Data analyzed using unpaired one-way ANOVA; data are mean ± s.d. with each dot representing individual mice. * *P* < 0.05, **** *P* < 0.0001.

LCs in the epidermis typically have a low proliferation rate under homeostasis, so we expected their numbers to increase slowly after wound closure ^8–10^. However, we detected a transient and dramatic increase in LCs in the re-epithelialized area at Day 15 following injury (Figure 3A-C). To monitor these changes in LC density, we recorded time-lapse movies. Although LC division is uncommon at homeostasis, by Day 7 after injury, we observed multiple LCs dividing in the re-epithelialized region (Figure 3D, E, and Video S3, 4). To accurately measure proliferation over time, we performed whole-mount staining for Ki67 at multiple post-injury time points. Quantitative analysis revealed a gradual increase in total Ki67-positive cells (mostly epithelial cells), which peaked at Day 13 (Figure 3F, G). However, proliferating LCs (double positive for Ki67 and MHCII) exhibited an earlier peak at Day 7 before declining by Day 13. (Figures 3F, H, and S3A, B). Because CSF1R signaling regulates LC maintenance and proliferation, we next measured CSF1 and IL-34 protein levels at the wound site^5,25^. IL-34, but not CSF1, was significantly increased at Day 7 and was enriched in the epidermal fraction, raising the possibility that epidermal IL-34 contributes to the local increase in LC proliferation after wound closure (Figure S4A-D).

Unexpectedly, the proliferation surge resulted in an LC density exceeding normal levels in the wound site at Day 15 (Figure 3C). To explore the mechanisms underlying the decline in LCs after Day 15 following injury, we quantified apoptotic cells by staining for cleaved-caspase 3. Three weeks after injury, we detected apoptotic cells, including LCs, in the re-epithelialized region, where the frequency of apoptotic LCs was higher than in homeostatic skin. (Figure 3I-K). Additionally, we observed unexpected LC shedding at the wound site. Some LCs migrated to the superficial layers of the epidermis and subsequently shed off, a process similar to epithelial cell shedding in cornified layers (Figure S5A-C)^26^. Despite the drastic changes in cell density, LCs eventually recovered their spatial organization, with an evenly spaced distribution across the epidermis (Figure S5D, E). We did not detect either apoptosis or shedding in normal, uninjured epidermal LCs, indicating that LC death and shedding during wound healing are rare or specialized phenomena specific to the repair context.

Taken together, these data suggest that LC density in the regenerated epidermis is restored through temporally coordinated phases of LC proliferation followed by LC loss.

## Two populations of LCs contribute to LC network restoration at the wound site

LCs in adult skin consist of embryonically-derived resident LCs (eLCs) that maintain homeostasis^8^. Upon substantial eLC loss, circulating progenitor cells, which often correspond to monocytes^14^, infiltrate the epidermis via hair follicles and differentiate into LCs about 2 weeks after LC depletion^11–14^. Given our observation that LC numbers increased at the wound site around 2 weeks after injury, we hypothesized that, in addition to eLCs, progenitor-derived LCs also contribute to LC repopulation.

To test this, we developed a dual-labeling mouse line (*huLangerin-CreER; Rosa-stop-tdTomato; Lang-EGFP; K14-H2B-mCherry*) that distinguishes newly differentiated LCs from pre-existing eLCs (Figures 4A and S2A). In this system, all LCs express cytoplasmic EGFP. Following tamoxifen injection, eLCs also express tdTomato and appear yellow, whereas progenitor-derived LCs that arise after the 24-48 h tamoxifen window express only EGFP and appear green ^27^.

**Figure 4.**
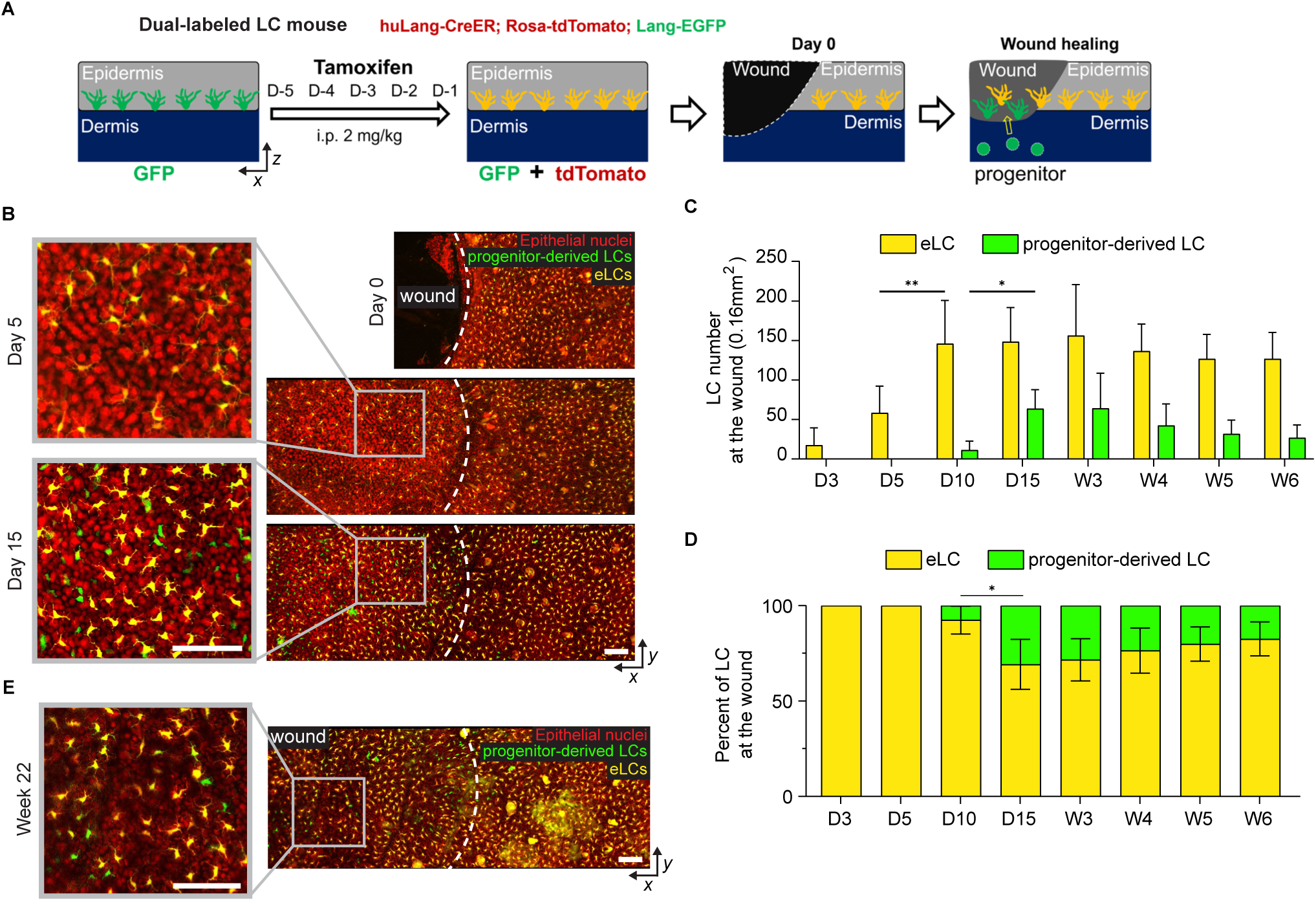
Multiple sources of Langerhans cells repopulate the wound site. **A**, Experimental design of dual-labeled LC mouse line. Dual-labeled LC mice received daily tamoxifen injections 5 days prior to wound induction. Embryonic LCs (eLCs) are labeled yellow and progenitor-derived LCs are only green. **B**, Revisit multi-photon *in-vivo* microscopy images of a 1 mm wound from the same mouse. Images show *x-y* view of epithelial cells (red nuclei), eLCs (yellow), and progenitor-derived LCs (green) in the epidermis at 5 and 15 days after wound induction. Dashed line indicates initial wound boundary. *Left*: zoomed view of the wound center. Representative images from 6 mice. Representative image from 3 mice. Scale bars, 100 µm. **C**, Timeline of LC number comparing cell density changes among embryonic and progenitor-derived LCs at the wound. *n* = 6 mice. **D**, Timeline of changes in the percentage ratio between embryonic and progenitor-derived LCs at the wound site. *n* = 6 mice. **C-D**, Wound area quantified 0.16 mm^2^ per mouse. Data analyzed using paired two-way ANOVA; data are mean ± s.d. * *P* < 0.05, ** *P* < 0.01. **E**, Multi-photon *in-vivo* microscopy of dual-labeled LC mice 22 weeks after wound induction. Dashed line indicates initial wound boundary. *Left*: zoomed view of the wound center. Representative image from 3 mice. Scale bars, 100 µm.

We administered tamoxifen to the skin prior to wounding, thus dual-labeling all eLCs. At Day 5 after injury, only yellow LCs were observed at the wound site (Figure 4B), confirming that resident eLCs migrate to the wound during re-epithelialization. As we hypothesized, green progenitor-derived LCs were present at the wound site on Day 15 post-injury (Figure 4B), indicating that this population also contributes to LC repopulation after injury^11–14^. Progenitor-derived LCs appeared smaller in volume and less dendritic than eLCs (Figure 4B), similar to monocyte-derived LCs reported in previous studies^12^. Nevertheless, both populations were evenly distributed in the re-epithelialized epidermis (Figure 4B), suggesting their integration into a unified LC network.

Notably, although LC density dropped at both the wound and the near regions (Figure 3A-C), progenitor-derived LCs repopulated primarily at the wound site (Figure 4B), implying that recruitment is triggered by wound-specific changes rather than a local drop in LC density. Quantification revealed that progenitor-derived LCs were rare on Day 10 after injury, peaked at Day 15-21, and then declined during the late remodeling phase (Figure 4C). In contrast, eLCs peaked at Day 10 after injury and did not decline substantially (Figure 4C). Indeed, the progenitor-derived LC contribution to LC density peaked at ∼30% at Day 15 and was ∼18% by week 6 post-injury (Figure 4D). These data suggest that eLCs are preferentially retained over progenitor-derived LCs as part of the LC network.

Whether progenitor-derived LCs are short-lived replacements or serve as a long-term, bona fide supplement to the resident LC network has remained unclear^12,13^. In the context of wound healing, we found that progenitor-derived LCs persisted in the re-epithelialized region at 22 weeks post-injury, suggesting that these cells are long-lived (Figure 4E). We infer that progenitor-derived LCs are not merely transient replacements but can establish themselves as long-term components of the LC network after injury.

Altogether, our results revealed that, despite differences in origin and morphology, eLCs and progenitor-derived LCs together reestablish the LC population at the wound site in the epidermis. In addition, although eLCs are preferentially maintained, progenitor-derived LCs persist in the epidermis long after wound healing.

## Monocytes recruited to the wound site differentiate into mLCs

Under inflammatory conditions, LC replenishment can involve monocytes as well as additional myeloid precursors^11–14,28^. To determine if progenitor-derived LCs are derived from monocytes after wounding, and if the two LC populations differ at a transcriptomic level, we performed single-cell RNA sequencing (scRNA-seq) on sorted LCs from tissue collected on Day 11 after injury. We chose Day 11 because we aimed to capture progenitor-derived LCs early in their differentiation.

The UMAP cluster projection revealed a strong enrichment of LCs along smaller clusters belonging to other cell types (Figure S6A). All the cells passed basic quality control filtering, and the clusters were annotated according to their top differentially expressed genes (DEGs) (Figure S6B). LC clusters were identified by high *Langerin* (*Cd207*) expression and divided into five subsets: activated (act. LC; high *Cd80*, low *Cdh1*), steady-state (s.s. LC; low *Cd80*, high *Cdh1*), cycling (cyc. LC; *Top2a*), migrating (mig. LC; *Ccr7*), and apoptotic (apop. LC; *Ddias*).

Importantly, we identified a cluster that showed moderate expression of the monocyte marker *Cd68* and the LC marker *Cd207*, which likely contained the newly differentiated progenitor-derived LCs (Figure S6C). To explore whether monocytes give rise to progenitor-derived LCs after skin injury, we subset our data to include only the monocyte and LC clusters. From the re-clustered data, we created a new UMAP and assigned cells to four main clusters: monocytes (Mono), pre-monocyte-derived LCs (pre-mLC), monocyte-derived LCs (mLC), and embryonic LCs (eLC) (Figure 5A). Across the four clusters, there was a gradual reduction of the monocytic marker *Cd14* going from the Mono to the eLC cluster. Notably, the mLC cluster showed higher *Cd14* expression than the eLC cluster. Conversely, there was an increase in *Cd207* expression from the Mono cluster to the eLC cluster (Figure 5B).

**Figure 5.**
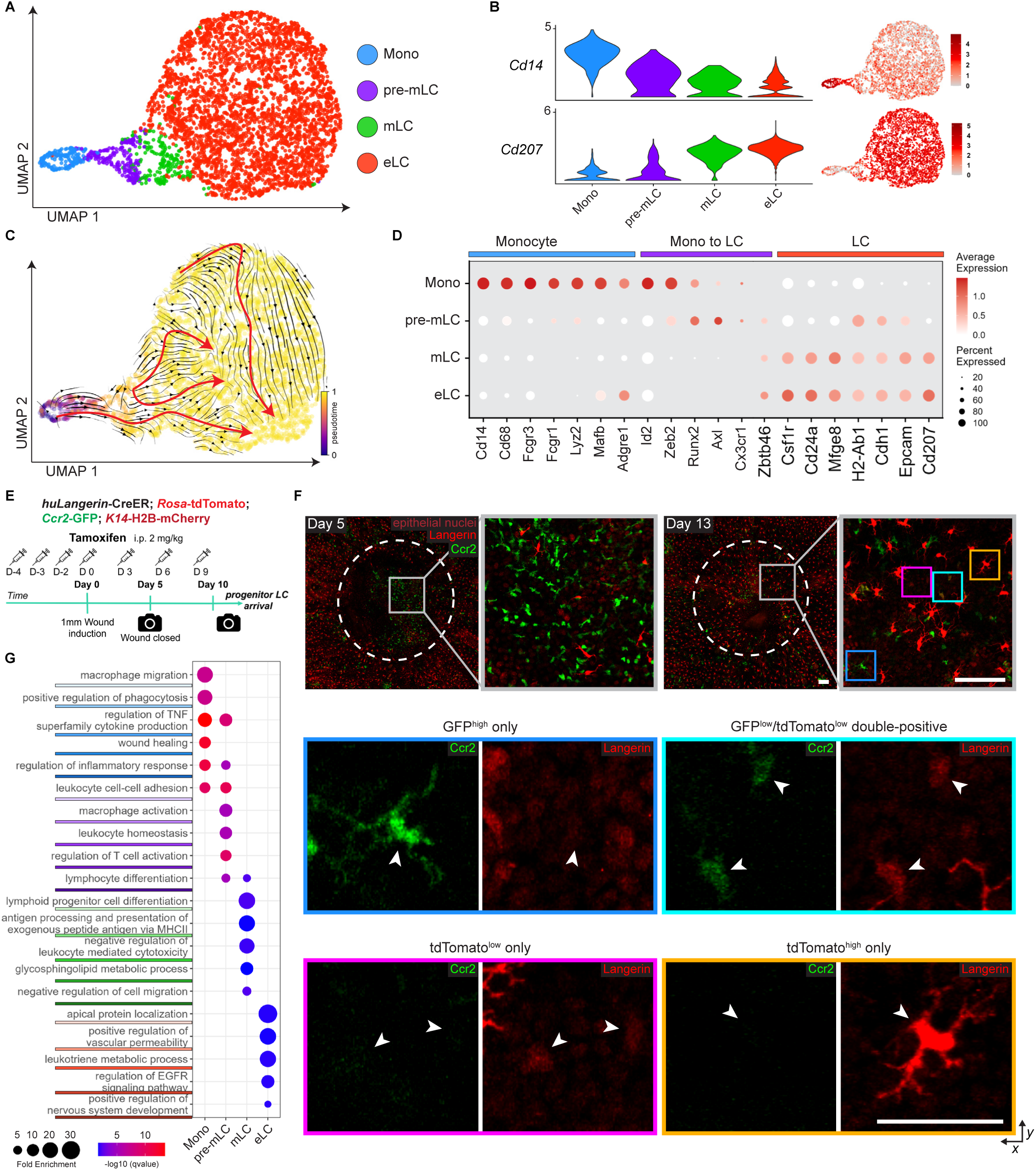
Monocyte-derived Langerhans cells integrate into the existing LC network after wound closure. **A**, UMAP of monocyte and Langerhans cells re-clustering. Clusters were defined as monocytes (Mono), pre-monocyte-derived LCs (pre-mLC), monocyte-derived LCs (mLC), and embryonic LCs (eLC). Embryonic LCs subclusters were merged as “eLC” for simplicity. **B**, *Left*: Violin plots showing the expression of top markers for monocytes (*Cd14*) and LCs (*Cd207*). *Right*: Feature plots showing the expression of the gene mentioned in the same row. **C**, RNA velocity pseudotime trajectories. The direction of the black arrows reflects the changes in the surrounding cellular states. Red arrows show the general trend. **D**, Dot plot of selected marker genes for monocytes, LC differentiation, and embryonic LCs. **E**, Experimental design for the *in-vivo* identification of repopulating monocyte-derived LCs. Tamoxifen was used to label existing embryonic LCs red prior to wound induction. Tamoxifen continued to be delivered up until the expected arrival of progenitor LCs (Day 10 after wound induction). Injections were delivered every 3 days after wound induction to avoid toxicity. Revisit imaging (camera icon) was performed on Day 5 and 13. **F**, Revisit multi-photon *in-vivo* microscopy images of a 1 mm wound from the same mouse. *Top*: Images show *x-y* view of epithelial cells (dim red nuclei), monocytes (green), and LCs (red) in the epidermis at 5 and 13 days after wound induction. *Blue frame*: zoomed example of a GFP^high^ only cell. *Teal frame*: zoomed example of GFP^low^/tdTomato^low^ double-positive cells. *Magenta frame*: zoomed example of tdTomato^low^ only cells. *Orange frame*: zoomed example of a tdTomato^high^ only cell. White arrowheads highlight the same cell within each frame. Dashed line indicates initial wound boundary. Representative images from 3 mice. Scale bars: *Top*: 100 µm, *Colored frames*: 50 µm. **G**, GO plot of selected biological process terms among the top 50 results enriched in each cluster using the top 200 markers. Highlighted color shades under GO terms match the corresponding cluster colors.

To further characterize monocyte-to-mLC differentiation, we performed RNA velocity analysis. Pseudotime trajectories showed directionality that suggests that cells from the Mono cluster pass through the pre-mLC cluster and eventually reach the mLC cluster (Figure 5C). We also found that within the eLC cluster, epidermal LCs transitioned from steady-state (low *Cd80*, high *Cdh1*) to activated (high *Cd80*, low *Cdh1*)^24,29^ (Figures 5C and S7A, B). Next, we compared the expression of known monocytic (*Cd14, Cd68, Fcgr3, Fcgr1, Lyz2, Mafb,* and *Adgre1*) and LC (*Csf1r, Cd24a, Mfge8, H2-Ab1, Cdh1, Epcam,* and *Cd207*) marker genes, as well as genes described to play a role in LC differentiation (*Id2, Zeb2, Runx2, Axl, Cx3cr1,* and *Zbtb46*)^7,14,30–33^. Indeed, we found that the Mono and eLC clusters had strong expression of their corresponding marker genes. Importantly, the pre-mLC cluster showed specific expression of LC differentiation markers and low expression of LC markers. We also noticed that the mLC and eLC clusters strongly resembled each other and expressed LC markers (Figure 5D). This analysis suggests that, during wound healing, monocytes differentiate into mLCs that express the major LC signature genes. To validate the monocyte origin of mLCs, we performed adoptive transfer of CD45.2 monocytes into CD45.1 host mice followed by 1-mm wound induction. At Day 14 post-injury, donor-derived CD45.2⁺ cells were detected in the wound epidermis and co-expressed the LC marker CD207, confirming that transferred monocytes can acquire LC identity during wound healing (Figure S8A-D). Although donor-derived cells were detected at low frequency, likely due to contribution from endogenous host cells, these results provide direct experimental support for monocyte-to-LC differentiation.

Given that transferred monocytes can become LCs in the wounded epidermis, we next used intravital imaging to define the temporal dynamics of this differentiation process at the wound site. We generated a new fluorescent mouse line that tagged LCs in red and monocytes in green (*huLangerin-CreER; Rosa-stop-tdTomato; Ccr2-GFP; K14-H2BmCherry*)^8,14,34,35^. We used tamoxifen to activate tdTomato in eLCs prior to wound induction and continued injections up to when we had observed the first appearances of progenitor-derived LCs (Day 10 post-injury) (Figure 5E). Through intravital microscopy, we followed the fate of *Ccr2-GFP* expressing cells and determined their differentiation into LCs by expression of tdTomato in response to tamoxifen.

We observed monocytes (GFP-positive, tdTomato-negative) at the wound site as early as Day 5 post-injury. By Day 13, the wound site contained distinct cell populations: GFP^high^ only, GFP^low^/tdTomato^low^ double-positive, tdTomato^low^ only, and tdTomato^high^ only cells (Figure 5F). Notably, we did not observe bright tdTomato cells that retained GFP, suggesting that CCR2 and Langerin are not co-expressed in the same cells during this transition. To further characterize the CCR2-GFP⁺ cells present in the wound epidermis at Day 13, we performed flow cytometric analysis. The results showed that CCR2-GFP⁺ myeloid cells were predominantly CD207⁺ across both Ly6C⁻ and Ly6C⁺ populations (Figure S9A, B). These findings suggest that CCR2-GFP⁺ cells remaining in the wound epidermis until Day 13 may have already transitioned toward an LC-like state. These results provide direct in vivo evidence that monocytes enter the wound as CCR2+ precursors and progressively differentiate into LCs through distinct intermediate states. Thus, these findings reveal the cellular trajectory of monocyte-to-LC differentiation at the wound site and underscore an unexpected heterogeneity of this process.

Under immune-pathological conditions, mLCs become transcriptionally and functionally similar to eLCs^12^. To elucidate potentially distinct contributions of mLCs and eLCs during wound repair, we used the top 200 DEGs from each cluster to determine the top 5 most relevant gene ontology (GO) terms (Figure 5G). The Mono cluster was characterized by a unique role in macrophage migration, positive regulation of phagocytosis, and wound healing processes (Figures 5G and S7C). Both Mono and pre-mLC clusters shared enrichment in pathways involved in TNF production and the inflammatory response, as well as genes mediating leukocyte cell-cell adhesion. The pre-mLC cluster further exhibited unique enrichment for macrophage activation, leukocyte homeostasis, and regulation of T cell activation (Figures 5G and S7D). Interestingly, the pre-mLC and mLC clusters shared enrichment of genes related to lymphocyte differentiation, while the mLC cluster alone showed unique enrichment for lymphoid progenitor differentiation, antigen presentation, glycosphingolipid metabolism, and negative regulation of leukocyte-mediated cytotoxicity and cell migration (Figures 5G and S7E). In contrast, the eLC cluster demonstrated distinct enrichment of genes associated with apical protein localization, positive regulation of vascular permeability, leukotriene metabolism, EGFR signaling, and nervous system development (Figures 5G and S7F).

Altogether, our data support a model in which monocytes infiltrate the wound site in the epidermis and subsequently differentiate into long-term mLCs. While differentiating mLCs progressively acquire the transcriptional profile of steady-state eLCs, subtle differences in gene expression hint at a role for eLCs in tissue regeneration.

## A non-canonical chemokine receptor, CXCR2, drives directional eLC migration into wounds

Our results established that resident eLCs are the first LCs to migrate into the wound during re-epithelialization. As part of their role as APCs, LCs migrate out of the epidermis through the chemokine receptor *Cxcr4*^36^. Once in the dermis, LCs rely on another chemokine receptor, *Ccr7*, to enter the lymphatic vessels and continue their journey towards the lymph nodes^37^. To determine if either of these chemokine receptors was also important for LCs to mobilize within the epidermis after injury, we generated two loss-of-function mouse models. Specifically, we engineered an LC-specific *Cxcr4* KO mouse line (*huLangerin-CreER; Cxcr4^−/−^; Lang-EGFP; K14-H2B-mCherry*, hereafter referred to as LC-Cxcr4 KO), and a *Ccr7* global KO mouse line (*Ccr7^−/−^; Lang-EGFP; K14-H2B-mCherry*, hereafter referred to as Ccr7 KO). For the LC-Cxcr4 KO mice, we floxed out *Cxcr4* via tamoxifen prior to wound induction. On Day 5 post-injury, LC density in LC-*Cxcr4* KO wounds showed a slight increase compared with controls, whereas *Ccr7* KO wounds showed a slight decrease (Figure S10A, B). Despite these minor changes, the data indicate that neither *Cxcr4* nor *Ccr7* is required for eLC migration toward the wound during re-epithelialization.

To explore new mechanisms of eLC migration, we searched for alternative chemokine receptors expressed in eLCs. Briefly, we isolated eLCs and epithelial cells from mouse epidermis during homeostasis and performed bulk RNA-seq. We identified chemokine receptors that were most strongly expressed in homeostatic eLCs compared to surrounding epithelial cells, including *Cxcr2* (Figure 6A).

**Figure 6.**
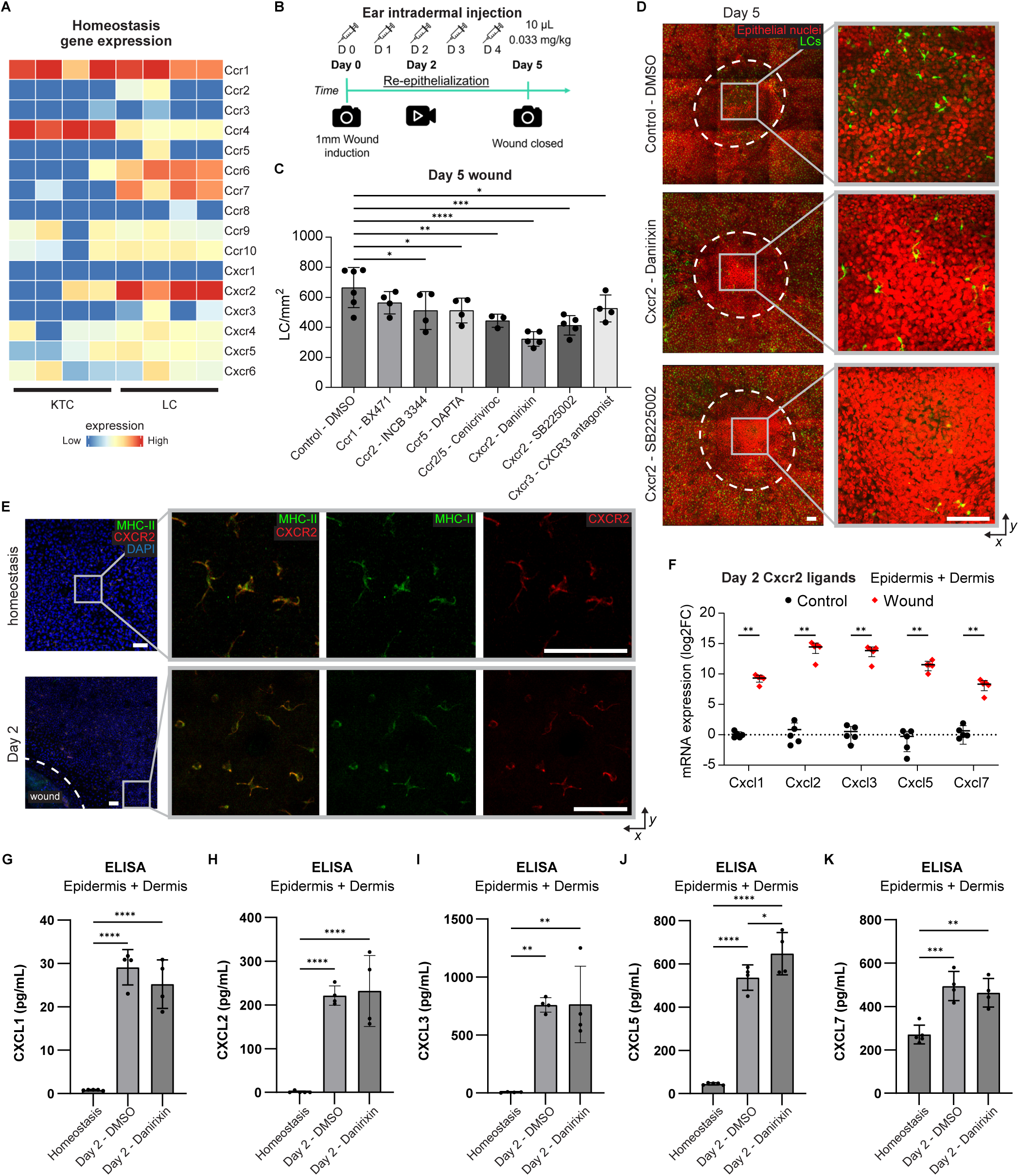
Cxcr2 inhibition blocks eLC wound repopulation. **A**, Heatmap of normalized log2 fold change of chemokine receptor gene expression from homeostatic epithelial (KTC) and Langerhans cells (LC). Cells isolated through FACS and sequenced through bulk RNA-seq. Each column represents an independent sample, and each row is assigned to a specific gene. Red indicates maximum expression and blue indicates minimum expression. *n* = 4 mice **B**, Experimental design for drug treatment. Starting on wound induction day, drug was injected once a day intradermally at the ear near the wound site. Control mice received vehicle (1% DMSO) injections. Wounds were imaged at wound closure (5 days after wound induction), and candidate drugs were further analyzed for migration dynamics through time-lapse at 2 days after wound induction. Revisit imaging (camera icon) was performed on Day 0 and 5. Timelapse imaging (video icon) was performed on Day 2. **C**, Mean LC number comparing cell density at the wound in response to drug treatment. Imaging was performed 5 days after wound induction. LC density normalized to the individual mouse wound area was quantified. Data analyzed using unpaired one-way ANOVA; n = 6 control, n = 4 BX471, n = 4 INCB3344, n = 4 DAPTA, n = 3 Cenicriviroc, n = 5 Danirixin, n = 5 SB225002, and n = 4 CXCR3 antagonist-treated mice.; data are mean ± s.d. with each dot representing individual mice. * *P* < 0.05, ** *P* < 0.01, *** *P* < 0.001, **** *P* < 0.0001. **D**, *In-vivo* microscopy images show *x-y* view of epithelial cells (red nuclei) and LCs (green) in the epidermis 5 days after wound induction. *Top*: control mouse (1% DMSO). *Middle*: CXCR2-inhibited mouse (Danirixin). *Bottom*: CXCR2-inhibited mouse (SB225002). *Right*: zoomed view of the wound center matching the image on the left. Dashed line indicates initial wound boundary. Representative images are shown. n = 6 control mice and n = 5 mice per drug-treated group. Scale bars, 100 µm. **E**, Confocal immunofluorescent images of CXCR2 expression at the epidermis during homeostasis and 2 days after wound induction. Images show *x-y* view of LCs (green, MHC-II), CXCR2 (red), and cell nuclei (blue, DAPI). *Right*: zoomed view in composite, green channel only, and red channel only. Dashed line indicates initial wound boundary. Representative images from 3 mice. Scale bars, 25 µm. **F**, qRT-PCR gene expression analysis of CXCR2 ligands in the skin during homeostasis (control) and 2 days after wound induction. Data analyzed using multiple unpaired two-tailed *t*-test; *n* = 5 mice; data are mean ± s.d. with each dot representing individual mice. ** *P* < 0.01. **G**, ELISA assay of CXCL1 present on wounds treated with CXCR2-inhibitor (Danirixin) compared to control (DMSO) and homeostasis. **H**, ELISA assay of CXCL2 present on wounds treated with CXCR2-inhibitor (Danirixin) compared to control (DMSO) and homeostasis. **I**, ELISA assay of CXCL3 present on wounds treated with CXCR2-inhibitor (Danirixin) compared to control (DMSO) and homeostasis. **J**, ELISA assay of CXCL5 present on wounds treated with CXCR2-inhibitor (Danirixin) compared to control (DMSO) and homeostasis. **K**, ELISA assay of CXCL7 present on wounds treated with CXCR2-inhibitor (Danirixin) compared to control (DMSO) and homeostasis. **G-K**, Data analyzed using unpaired one-way ANOVA; *n = 4* mice; data are mean ± s.d. with each dot representing individual mice. * *P* < 0.05, ** *P* < 0.01, *** *P* < 0.001, **** *P* < 0.0001.

To test the role of these receptors in eLC migration during re-epithelialization after injury, we designed a chemokine receptor drug inhibitor screen (Figure 6B). We targeted *Ccr1* and *Cxcr2*, as well as *Ccr2*, *Ccr5*, and *Cxcr3*, based on the availability of drug antagonists. We excluded *Ccr7* and *Cxcr4* since we had already tested them using functional mouse models. Relative to control (vehicle), almost all chemokine receptor inhibitors resulted in a modest decrease in eLC density at the wound site on Day 5 post-injury, with two different CXCR2 inhibitors resulting in the largest decreases (Figure 6C, D). Using whole-mount immunofluorescent staining, we confirmed that CXCR2 protein is specifically expressed by eLCs, not by epithelial cells, in the epidermis during homeostasis and after injury (Figure 6E). To corroborate the function of CXCR2, we assessed the expression of CXCR2 ligands at the wound site during re-epithelialization. We found a significant increase in the expression of all CXCR2 ligands (*Cxcl1*, *Cxcl2*, *Cxcl3*, *Cxcl5*, and *Cxcl7*) at both the RNA and protein levels on Day 2 after injury compared to homeostasis (Figure 6F-K). Notably, Danirixin treatment did not substantially change the protein levels of these ligands compared with DMSO-treated wounds, suggesting that CXCR2 blockade does not broadly affect ligand expression at the wound site.

Taken together, these results suggest that LCs employ a novel pathway for migration into wounds during re-epithelialization via CXCR2–ligand signaling, highlighting a non-canonical mechanism distinct from the pathways that guide LC egress to lymph nodes.

## CXCR2 inhibition redirects wound repopulation from eLCs to mLCs

We have shown that eLCs migrate towards the wound (Figure 1C-E) and that inhibition of CXCR2 decreases LC density at the wound site (Figure 6C, D). Based on these data, we hypothesized that CXCR2 regulates the movement of LCs into the wound epidermis during re-epithelialization. To test this hypothesis, we used time-lapse imaging to investigate the effect of CXCR2 inhibition on the real-time behavior of migrating LCs on Day 2 after injury (Figure 7A, B and Video S5). First, we evaluated whether CXCR2 inhibition affected epithelial cell migration and found that epithelial cell displacement showed no significant difference between drug-treated and control mice (Figure 7C). Notably, inhibition of Cxcr2 did not cause any delay in wound healing (Figure 6D). Next, we examined LC track displacement. CXCR2 inhibition did not significantly reduce LC mobility (Figure 7D). However, when we analyzed LC directionality, we found that LCs failed to mobilize towards the wound in mice treated with CXCR2 inhibitor compared to control-treated mice (Figure 7E). Thus, wounding is sufficient to trigger LC mobility, but *Cxcr2* is required to direct LC movement towards the wound during re-epithelialization.

**Figure 7.**
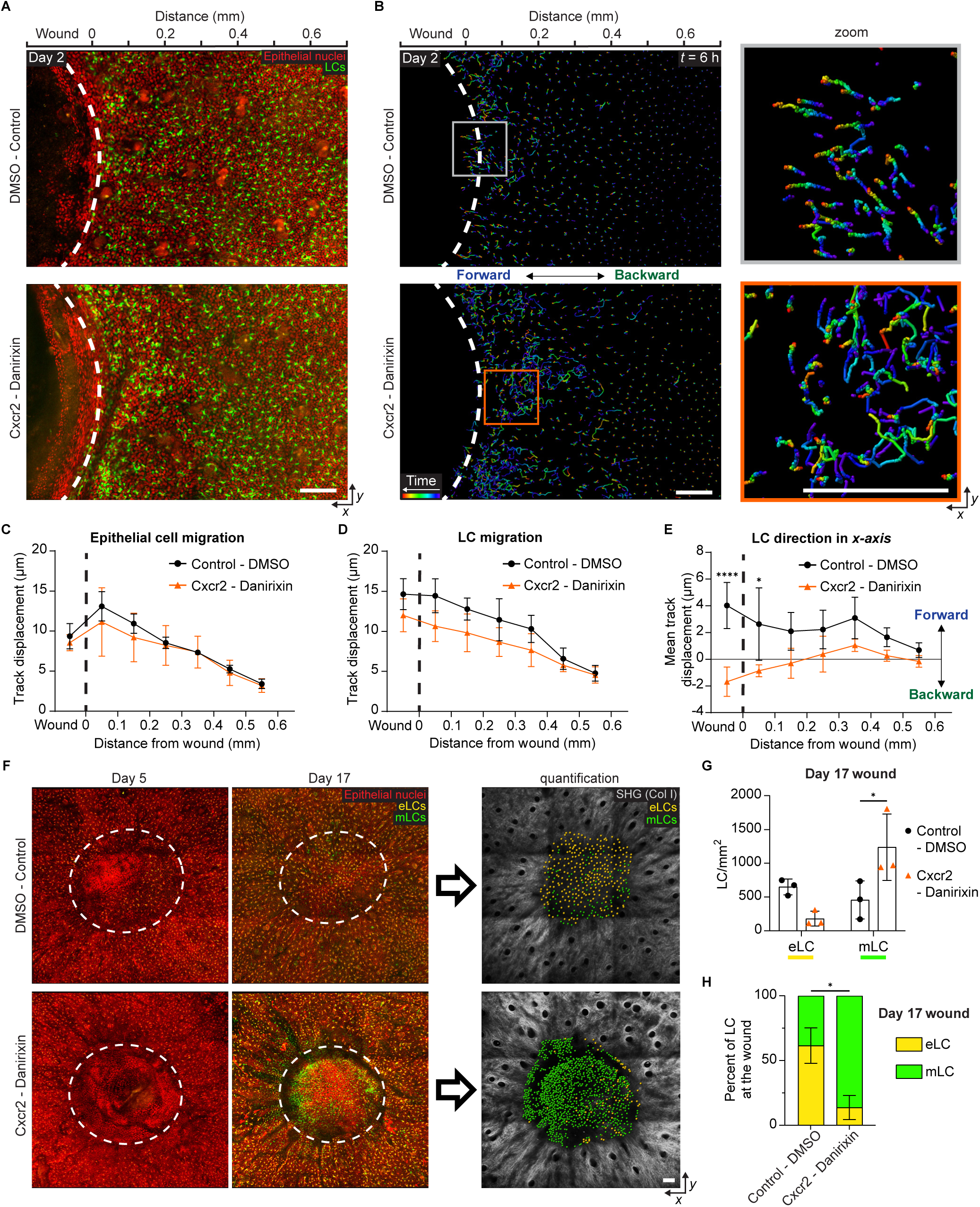
Monocyte-derived Langerhans cells compensate for the loss of embryonic Langerhans cells at the wound. **A**, Time-lapse *x-y* view of epithelial cells (red nuclei) and LCs (green) 2 days after wound induction. *Top*: control mouse 1% DMSO. *Bottom*: drug-treated mouse CXCR2 (Danirixin) as shown in (**Figure 6B**). Dashed line indicates initial wound boundary. Representative images from 3 mice per group. Scale bars, 100 µm. **B**, Imaris *x-y* view track analysis of LCs (**A**) 2 days after wound induction. Colors project time (blue, 0h; red, 6h). *Top*: control mouse 1% DMSO. *Bottom*: drug-treated mouse CXCR2 (Danirixin). *Right*: zoomed migration tracks from near the wound edge. Representative images from 3 mice per group. **C**, Mean total displacement of individual epithelial cells tracks from control and CXCR2-inhibited mice over 6h plotted as a function of distance from the wound. *n* = 3 mice. **D**, Mean total displacement of individual LC tracks from control and CXCR2-inhibited mice over 6h plotted as a function of distance from the wound. *n* = 3 mice. **E**, Mean track displacement in the *x* axis of individual LC tracks from control and CXCR2-inhibited mice over 6h plotted as a function of distance from the wound. Calculated by comparing the start and end values on the *x* axis of each track. Positive change indicates movement towards the wound. *n* = 3 mice. **C-E**, Imaging performed 2 days after wound induction. Dashed line, initial wound boundary. The displacements of migrating cell tracks were averaged every 100 µm from the initial wound. Data analyzed using unpaired two-way ANOVA; data are mean ± s.d. * *P* < 0.05, **** *P* < 0.0001. **F**, Revisit multi-photon *in-vivo* microscopy images of a 1 mm wound from the same mouse. Images show *x-y* view of epithelial cells (dim red nuclei), embryonic LCs (orange/yellow), and progenitor-derived LCs (green) in the epidermis at 5 (*Left*) and 17 (*Middle*) days after wound induction. Day 17 LCs at the wound quantified using Imaris spots analysis (*Right*). *Top*: control mouse 1% DMSO. *Bottom*: drug-treated mouse CXCR2 (Danirixin). Dashed line indicates initial wound boundary. Representative images from 3 mice per group. Scale bar, 100 µm. **G**, Mean LC number comparing cell density changes among embryonic LCs (eLC) and monocyte-derived LCs (mLC) in response to CXCR2 inhibition. **H**, Percentage ratio between eLCs and mLCs at the wound epidermis in response to CXCR2 inhibition. **G-H**, Imaging performed 17 days after wound induction. LC density normalized to the individual mouse wound area was quantified. Data analyzed using unpaired two-way ANOVA; *n*≥3 mice; data are mean ± s.d. * *P* < 0.05.

Given that eLCs and mLCs both repopulate the wound epidermis, we wondered if blocking the directional migration of eLCs would affect LC repopulation dynamics during healing. To test this, we used our approach above to inhibit CXCR2 in our dual-label LC mouse line that distinguishes eLCs from mLCs (Figure 4A, 6B). We tracked the changes of the two LC populations in CXCR2-inhibited and control mice on Days 5 and 17 after injury using in vivo microscopy. Imaging on Day 5 confirmed a reduction in eLCs within the wound closure site in drug-treated mice (Figure 7F). Quantification on Day 17 showed a decrease in eLC density and a concomitant increase in recruited mLCs in mice treated with CXCR2 inhibitor compared to control-treated mice (Figure 7F, G). Inhibition of CXCR2 caused a substantial increase in the ratio of mLCs to eLCs at the wound closure site (Figure 7H). The increase in mLCs accompanied by the decrease in eLCs, along with the maintenance of total LC density, suggest functional compensation.

Altogether, the data point to a versatile LC repopulation, where the loss of eLCs at the wound epidermis is rebalanced by a higher repopulation of mLCs, suggesting that mLCs compensate for eLC deficiency to maintain the LC pool.

## Discussion

LCs function as first responders in skin immunity, and their maintenance in the epidermis is of utmost importance. Here, through intravital microscopy to longitudinally follow the fate of LCs, we find that recovery of the lost niche after wound repair involves the coordinated repopulation by two distinct LC lineages. During re-epithelialization, eLCs migrate intraepidermally into the wound, and after closure, the LC pool expands through local proliferation within the regenerated epidermis. In parallel, circulating monocytes infiltrate the wound and differentiate in situ into long-lived mLCs, contributing to durable network reconstitution. Together, these processes restore LC density and spatial organization, a structural benchmark for re-establishing the epidermal immune barrier.

LCs are widely recognized for their function as APCs. This role requires antigen capture and migration out of the epidermis, towards the lymph nodes. Our time-lapse and revisit imaging show that LCs adjacent to the wound actively migrate into the wound bed during re-epithelialization. Consistent with a wound-directed program, genetic ablation of the lymphatic-homing receptors Cxcr4 or Ccr7 did not substantially alter LC numbers at the re-epithelialized site, suggesting that only a small fraction of LCs leave toward the dermis during wound repair, while the majority remain in the epidermis.

The migration of epidermal-resident LCs into the wound bed is temporally aligned with epithelial cell migration. Our data show that injury stimulates cell-autonomous LC mobility adjacent to the wound independently of epithelial cell movement, as demonstrated through the selective disruption of *Rac1* signaling in either epithelial cells or LCs. Notably, we did not detect proliferation in actively migrating LCs during re-epithelialization; instead, a wound-specific hyperproliferation became evident immediately after closure. A similar event has also been observed during the first week of murine life, as well as under MC903-induced skin inflammation^38^. It has been proposed that an unknown signal released by epithelial cells drives LC proliferation^38^.

In addition to LC expansion through proliferation and repopulation, our time-lapse imaging revealed that LC removal from the epidermal network can occur through multiple mechanisms. Beyond apoptosis, we also observed shedding as an alternative route of LC elimination during wound repair. This raises the possibility that, analogous to the complementary mechanisms of LC expansion through proliferation and progenitor-derived repopulation, LC removal from the epidermal network may also be achieved through more than one pathway. Although both events were reproducibly captured, they involved only a small number of cells, and the mechanisms governing how individual LCs are selected for each fate remain unclear. Future studies will be needed to elucidate the signals that direct LCs toward apoptosis or shedding during tissue repair.

A severe loss of LCs in the uninjured state, induced by UV irradiation or immune-mediated pathology, has shown that recovery of the LC population arises from the differentiation of recruited monocytes via epithelial cells in hair follicles^11–14^. In the context of wound repair, we find that monocytes infiltrate the wound epidermis, which lacks hair follicles, by about Day 5 and differentiate into mLCs about two weeks post-injury. Notably, monocyte differentiation occurs within the wound epidermis only, suggesting that it is driven by wound-specific changes and is not restricted to hair follicles^7,14^. Furthermore, long-term longitudinal imaging demonstrates that mLCs can survive in the healed epidermis for up to 22 weeks.

Transcriptomic analyses provide an unbiased, high-resolution comparison of the two lineages during repair: mLCs acquire a profile reminiscent of steady-state LCs with tolerogenic programs, whereas eLCs are enriched for pathways linked to tissue regeneration (including nervous system development). These data support complementary roles for the two lineages in wound healing. Although our transcriptional and imaging data suggest complementary roles for eLCs and mLCs during wound repair, direct functional comparisons between these two LC populations would help clarify whether they have overlapping or distinct functions. Future lineage-specific perturbation studies may reveal whether eLCs and mLCs differ in antigen presentation, immune regulation, tissue remodeling, or other wound-associated functions. The participation of LCs in wound repair has been widely debated, with reports that they promote or hinder epithelial wound closure^39,40^. A recent study has demonstrated that LCs are essential contributors to angiogenesis during wound repair^41^. Together with our data, this could suggest a special role for eLCs in rebuilding skin architecture rather than wound closure. A similar ability by mLCs remains to be tested.

Finally, we identify the chemokine receptor *Cxcr2* as a critical determinant of eLC intraepidermal migration to the regenerating epidermis. Pharmacological inhibition of CXCR2 significantly impaired the infiltration of eLCs into the wound, suggesting that *Cxcr2* signaling is required for LC homing to injured epidermis. Indeed, we found that CXCL family chemokines (e.g., CXCL1, CXCL2, CXCL5) were upregulated in the wound milieu. These findings reveal a previously uncharacterized mechanism of intraepidermal LC guidance, distinct from the CXCR4-CCR7-mediated migration to lymph nodes classically described during immune activation. In addition to blocking eLC wound repopulation, pharmacological inhibition of CXCR2 increased mLC presence at the wound closure site, thereby maintaining LC density inside the wound. Because monocytes can also express CXCR2, lineage-specific genetic tests will be valuable to sharpen cellular specificity. Nevertheless, the inverse eLC–mLC relationship observed under CXCR2 blockade supports a compensatory repopulation dynamic that preserves the immune barrier.

Taken together, these findings suggest that LCs are not only immune sentinels for adaptive immunity but also dynamic responders to epidermal injury. By delineating the migratory dynamics, ontogeny-specific contributions, and chemokine responsiveness of LCs during wound healing, this study outlines a coordinated, two-lineage mechanism that rebuilds the epidermal LC compartment following injury. Our findings lay the groundwork for a more comprehensive understanding of LC biology during epidermal regeneration.

## Methods

### Mice and experimental conditions

*huLangerin-CreER*^42^ (JAX 028287), *Rosa-stop-tdTomato*^43^ (JAX 009669), *Ccr2-GFP*^44^ (JAX 027619), *Cx3cr1-GFP*^31^ (JAX 005582), *K14-CreER*^45^ (JAX 005107), *Rac1^fl/fl^*^46^ (JAX 005550), *Cxcr4 ^fl/fl^*^47^ (JAX 008767) and *Ccr7 KO*^48^ (JAX 006621) mice were obtained from Jackson Laboratories. *Lang-EGFP*^21^, *K14-H2B-mCherry*^22^, and *K14-H2B-Cerulean* ^20^ mice were obtained from V. Greco (Yale University). To simultaneously visualize LCs and epithelial cells, *Lang-EGFP*; *K14-H2B-mCherry* mice were generated. To knockout *Rac1* in epithelial cells, *K14-CreER*; *Rac1^−/−^* mice were mated with *Lang-EGFP*; *K14-H2B-mCherry* mice (*K14-CreER*; *Rac1^−/−^*; *Lang-EGFP*; *K14-H2B-mCherry*), and these mice were given five intraperitoneal injections of tamoxifen prior to wound induction (2 mg in corn oil per day for 5 days). Heterozygous siblings (*K14-CreER*, *Rac1^+/−^*, *Lang-EGFP*, and *K14-H2B-mCherry*) were used as controls. Similarly, to knockout *Rac1* in LCs, *huLangerin-CreER*; *Rac1^−/−^* mice were mated with *Lang-EGFP; K14-H2B-mCherry* mice (*huLangerin-CreER*; *Rac1^−/−^*; *Lang-EGFP*; *K14-H2B-mCherry*), and these mice were given five intraperitoneal injections of tamoxifen prior to wound induction (2 mg in corn oil per day for 5 days). Wild-type siblings were used as controls (*huLangerin-CreER*; *Rac1^+/+^*; *Lang-EGFP; K14-H2B-mCherry*). To distinguish between embryonic and progenitor-derived LCs, *huLangerin-CreER*; *Rosa-stop-tdTomato* mice were mated with *Lang-EGFP*; *K14-H2B-mCherry* mice (*huLangerin-CreER*; *Rosa-stop-tdTomato*; *Lang-\EGFP*; *K14-H2B-mCherry*), and these mice were given five intraperitoneal injections of tamoxifen prior to wound induction (2 mg in corn oil per day for 5 days). These mice were also used for scRNA-seq. To perform monocyte adoptive transfer assays, the bone marrow cells of wildtype B6 (CD45.2, JAX 000664) donor mice were harvested, and MACS enriched for monocytes. These monocytes were subsequently delivered via intravenous tail injection into a JAXBoy (CD45.1, JAX 033076) host mouse. To track monocytes recruited to the wound epidermis, *Ccr2-GFP* mice were mated with *huLangerin-CreER*; *Rosa-stop-tdTomato*; *K14-H2B-mCherry* mice (*huLangerin-CreER*; *Rosa-stop-tdTomato*; *Ccr2-GFP ^+/−^*; *K14-H2B-mCherry*), and these mice were given five intraperitoneal injections of tamoxifen prior to wound induction (2 mg in corn oil per day for 5 days). To label monocyte-derived LCs, these mice received four additional intraperitoneal injections of tamoxifen after wound induction (2 mg in corn oil on Days 5, 7, 9, and 11 post-injury). To isolate epithelial cells and LCs for bulk RNA-seq, we used a fluorescent mouse line established in our previous study. (*huLangerin-CreER*; *Rosa-stop-tdTomato*; *Cx3cr1-GFP ^+/−^*; *K14-H2B-Cerulean*)^20^. These mice labeled LCs in red, DETCs in green, and epithelial cells in blue. Wild-type CD1 mice were used for whole-mount staining and qRT-PCR experiments. Mice were housed on a ventilated rack with an ambient temperature of 22°C and a 12:12 light: dark cycle. Mice of both sexes from experimental and control groups were randomly selected for live imaging experiments. All mice used in this experiment were between 3 and 8 weeks of age. No blinding was done. All procedures involving animals were performed under the approval of the Institutional Animal Care and Use Committee (IACUC) at Michigan State University.

### Wounding

Three- to eight-week-old mice were anesthetized using an isoflurane chamber, and anesthesia was maintained throughout the course of the surgery with vaporized isoflurane delivered via a nose cone. Once a surgical plane of anesthesia was verified by the absence of a physical and psychological response to a noxious stimulus, a punch biopsy was performed using a 1-mm-diameter punch biopsy tool (Miltex). The punch biopsy tool was used to make a circular full-thickness wound on the dorsal side of the mouse ear. Wounds did not penetrate the ear and remained above the cartilage.

### Chemokine receptor inhibition

The following drug inhibitors were selected to target candidate LC chemokine receptors: BX-471 (Ccr1 antagonist, Cayman Chem.), INCB-3344 (Ccr2 antagonist, Cayman Chem.), DAPTA (Ccr5 antagonist, Cayman Chem.), Cenicriviroc (Ccr2/5 antagonist, Cayman Chem.), Danirixin (Cxcr2 antagonist, Cayman Chem.), SB-225002 (Cxcr2 antagonist, Cayman Chem.), and Cxcr3 antagonist 6c (Cxcr3 antagonist, Cayman Chem.). Drug inhibitors were dissolved in 1% Dimethyl sulfoxide (DMSO) in PBS to varied final concentrations (BX-471, 230 µM; INCB-3344, 173 µM; DAPTA, 117 µM; Cenicriviroc, 143 µM; Danirixin, 226 µM; SB-225002, 284 µM; Cxcr3 antagonist 6c, 162 µM). Control mice received only vehicle treatment (1% DMSO in PBS). A 10 µL dose (0.033 mg/kg) was intradermally injected at the ear wound site of *Lang-EGFP*; *K14-H2B-mCherry* daily for 5 days (Days 0, 1, 2, 3, and 4 after wound induction). To prevent disruption of the wound-healing process, the needle entry point was about 5 mm away from the wound site, and the dosage was administered slowly.

For these intradermal injection experiments, we used 8-week-old mice because their ear tissue was sufficiently large to allow stable delivery of 10 μL of solution. On Day 5, mice wounds were imaged and quantified for LCs located inside the wound epidermis. Only wounds completely closed by Day 5 were analyzed.

### Whole-mount staining

Ear tissues were processed for whole-mount staining. In brief, ears were incubated epidermis side up in 2.5% Trypsin solution at 37°C for 30 min and epidermis was removed from the dermis. The epidermis was fixed in 4% paraformaldehyde in PBS for 10 min at room temperature, washed in PBS, and then incubated in 20% sucrose in PBS at 4°C overnight. The samples were briefly washed in PBS and then blocked in 0.2% Triton X-100, 5% normal donkey serum, 1% BSA in PBS. The samples were then incubated with primary antibodies at 4°C overnight. Primary antibodies used were as follows: purified rat anti-mouse MHC class II (1:300, M5/114.15.2, Biolegend), purified rabbit anti-mouse Ki67 (1:400, D3B5, Cell Signaling Tech.), purified rabbit anti-mouse cleaved caspase-3 (Asp175) (5A1E) (1:300, 5A1E, Cell Signaling Tech.), AF647 CD207 (1:100, BDBiosciences; 564937). Adoptive transfer samples were stained with AF647 CD45.2 (1:100, Biolegend; 109818) and AF488 CD207 (1:100, Invitrogen; 53-2073-82). Samples were washed three times in 0.2% Triton in PBS for 1h each. The samples were then incubated with secondary antibodies at 4°C overnight. Secondary antibodies used were as follows: goat anti-rat Alexa Fluor 488 (1:300, Invitrogen), goat anti-rat Alexa Fluor 568 (1:300, Invitrogen), goat anti-rabbit Alexa Fluor 568 (1:300, Invitrogen). The tissue was also incubated with DAPI (1:1000) (Sigma, MBD0015) during secondary antibody exposure. Finally, samples were washed three times in 0.2% Triton in PBS for 1h each before mounting on glass slides. A STELLARIS 5 Leica confocal laser microscope was used to collect images.

### In vivo imaging

Mice were anesthetized using an isoflurane chamber and anesthesia was maintained throughout the course of the experiment with vaporized isoflurane delivered by a nose cone^18^. Mice were placed on a warming pad and kept at 38°C during imaging. The ear was mounted on a custom-made stage, and a glass coverslip was placed directly against it. Image stacks were acquired using a Leica SP8 DIVE laser multiphoton microscope equipped with Spectra-Physics Insight X3 dual beam (630 to 1300 nm tunable and 1040 nm fixed) and 4Tune, tunable, super-sensitive hybrid detectors (HyDs). Images were collected using Leica Application Suite X (LAS X) software (version 3.5.7.23225).

To acquire serial optical sections, a laser beam (940 nm) was focused through a ×25 water-immersion lens (NA 1.00 HC PL IRAPO, Leica) and scanned with a field of view of 0.59 × 0.59 mm^2^ at 600 Hz. To visualize a larger area, 2-25 tiles of optical fields were imaged using a motorized stage to automatically acquire sequential fields of view. Z stacks were acquired in 2 µm steps to image a total depth of 120 µm of tissue. Visualization of collagen was achieved via the second harmonic signal using the blue channel at 940 nm. For time-lapse imaging, serial optical sections were obtained in 4 min intervals. The duration of time-lapse imaging was 6-12h.

### Image analysis

Raw image stacks were imported into Fiji software (v1.53t; National Institute of Health) for tile merging and later analyzed using Imaris software (v10.0.0; Bitplane/Oxford Instruments). The tiled images were stitched by a grid/collection stitching plugin in Fiji. The merged image stacks were then imported into Imaris for further processing and analysis. LCs were identified and quantified (total cell number and average distance to the 3 nearest neighbors) from images using the Imaris Spots function, followed by individual examination for quality control (including cells not detected and excluding autofluorescence signals). To establish the LC distance from the wound, an artificial surface was created from the wound (identified as the area without second-harmonic generation), and the minimum distance of each LC spot to the surface was calculated. To quantify the LC distribution across the epidermal layers, an artificial surface was created using second-harmonic generation from fibrillar collagens. This surface roughly defines the basement membrane, in which the epithelial basal layer sits. The minimum distance between each LC spot and the basement membrane surface was then calculated.

For time-lapse videos, Imaris was used to track cells and obtain *xyz* coordinates from individual cells over time. Quality control was applied by individually examining all cell tracks to confirm that they reported the behavior of a single cell, as well as excluding tracks from dead cells from the skin surface and debris autofluorescence. Videos were corrected for drifting in the z-axis by applying Z-correction. To maintain consistency in video duration, only 6 hours were analyzed from each video. LC and epithelial cell migration were quantified by using the total length displacement from each track and by calculating their starting location in the x-axis in relation to the wound edge (identified by the area without second-harmonic generation). To determine the direction of LC migration towards the wound, the x-axis start value from each track was subtracted from the average change in the x-axis for that same track during the 6h analysis. Finally, individual cell tracks were binned every 100 µm from the wound, and the bin calculated means were used to yield trend lines.

### qRT-PCR

To quantify *Cxcr2* ligands (*Cxcl1*, *Cxcl2*, *Cxcl3*, *Cxcl5*, and *Cxcl7*), six-2mm punch biopsies were performed (as described in the wounding section) in a single mouse (3 per ear) with about 5 mm distance between them. On Day 2, a 3-mm punch biopsy was used to collect all six wounds, which included the epidermis and dermal layers of the dorsal side of the ear. Control samples were collected similarly from non-wounded mice. Total RNA was extracted using the Qiagen RNeasy Mini Kit, following the manufacturer’s instructions. RNA concentration was measured using the Qubit 4 Fluorometer with the RNA High Sensitivity Assay Kit (Thermo Fisher Scientific). Samples were diluted to a concentration of 200 ng/µL. cDNA was synthesized using the Qiagen QuantiTect Reverse Transcription Kit. Primers for qRT-PCR were designed using the Primer-BLAST tool. qRT-PCR reactions were performed using the Qiagen QuantiTect SYBR Green PCR Kit on a QuantStudio 5 Real-Time PCR System (Applied Biosystems). Gene expression levels were normalized to *Sdha*, and fold change values were calculated using the 2^−ΔΔCT method.

### Single-cell RNA sequencing sample preparation and data analysis

To isolate both progenitor-derived and embryonic LCs, we collected and pooled the ear epidermis of six *huLangerin-CreER*; *Rosa-stop-tdTomato*; *Lang-EGFP*; *K14-H2B-mCherry* mice into a single sample. These mice were all male littermates. To increase the collection of progenitor-derived LCs from the epidermis, wound induction was performed by softly applying microneedles (Skinmedix, micro needle dermal stamp 0.2 mm) repeatedly (approximately 500 times) across the dorsal side of the ear. This methodology increased the surface area of the damage applied, and consequently, a higher recruitment of progenitor-derived LCs at the epidermis. Ears were collected on Day 11 after wound induction. The ears were incubated epidermis side up in 2.5% Trypsin solution at 37°C for 90 min, and the epidermis was removed from the dermis.

Single cell suspension was achieved by mincing, cutting, and running the sample through a 70 µm filter. Samples were then enriched for LCs by magnetic sorting following the standard protocol from the Epidermal Langerhans Cell MicroBead Kit (Miltenyi Biotec). Upon inquiry, the manufacturer confirmed the potential off-target selection of other immune cell types.

Libraries were prepared using the 10x Chromium Next GEM Single Cell 3’ Kit, v3.1 and associated components. Libraries were pooled in equimolar proportions, then loaded onto 1 lane of an Illumina NovaSeq 6000 S4 flow cell (v1.5). The sequencing was performed in a 2x150bp paired-end format using a 300 NovaSeq v1.5 reagent cartridge. The fastq files were processed using the standard protocol that includes introns and exons from 10x Genomics Cell Ranger count (v8.0.0) with a custom Ensembl Mouse GRCm39 genome as the reference. The customized genome reference included a 2023 bp sequence, spanning from the tdTomato sequence to the downstream WPRE cassette ^49^, and a 1429 bp sequence which was assembled according to the original creation of the reporter mouse line EGFP ^43^. This custom sequence was incorporated into the reference genome following 10x Genomics guidelines. The data were then converted into a Seurat^50^ (v5.2.0) object.

The quality filtering was performed using the following criteria: nFeature_RNA: 3000–7000, nCount_RNA: 3000–70,000, percent_mito: <4.5%. To exclude keratinocytes, T cells, and neutrophils, the data were further filtered by normalized expression of the following genes: *Krt14*: <4, *Esrp1*: <1, *Cd3e*: <1, *S100a9*: <2. The data was scored for cell cycling genes and assigned an “S.Score” and “G2M.Score”. SCTransform^51^ was used to normalize and scale regress the data using (S.Score, G2M.Score, and percent_mito). The Louvain algorithm (Seurat-integrated) was used to generate the UMAP using the first 4 dimensions and a 0.6 resolution. Based on known gene markers, clusters were reassigned as follows: Monocytes (Mono), pre-monocyte-derived LCs (pre-mLC), monocyte-derived LCs (mLC), and embryonic LCs (eLC). Differential expression gene (DEG) analysis was performed using the built-in FindAllMarkers function in Seurat. The top 200 DEGs from each cluster were used to perform pathway enrichment analysis using the clusterProfiler^52^ package and determine the top 50 gene ontologies. For mRNA velocity analysis, Velocyto^53^ was used to calculate the spliced and unspliced read counts. The generated loom file was then merged with the UMAP coordinates and clusters into a Scanpy^54^ AnnData object. Secondary analyses and visualizations were performed using scVelo^55^.

### Bulk RNA sequencing sample preparation and data analysis

Single cell suspension was collected from four *huLangerin-CreER*; *Rosa-stop-tdTomato*; *Cx3cr1-GFP^+/-^*; *K14-H2B-Cerulean* mice during homeostasis following the same procedure as mentioned in the scRNA-seq section. LCs and epithelial cells were sorted using fluorescence-activated cell sorting (FACS) based on their red (LCs) and blue (epithelial cells) fluorescence. Libraries were prepared using the KAPA mRNA HyperPrep Kit with dUTP-based strand specificity. Libraries were normalized to 10 nM, then loaded onto an Illumina NovaSeq flow cell. The sequencing was performed using 100 bp paired-end sequencing on an Illumina HiSeq NovaSeq. Reads were aligned to the Ensembl Mouse GRCm39 genome using STAR Aligner^56^. DESeq2^57^ was used to determine the differential expression of chemokine receptor genes among the epithelial and Langerhans cell samples.

### Monocyte adoptive transfer

Bone marrow cells were pooled from both femurs and tibias from 3 donor CD45.2 mice. A 25-gauge syringe containing MACS buffer (1:20 dilution of MACS BSA Stock Solution with autoMACS Rinsing Solution) was used to flush the cells out. Cells were then filtered twice using a 70 µm and a 45 µm strainer, consecutively. Cells were then separated by following the standard procedure of the mouse BM MACS Monocyte Isolation kit (Miltenyibiotic; 130-100-629). Cell viability was assessed using trypan blue and monocyte enrichment was confirmed through flow cytometry. Following monocyte enrichment, cells were pelleted and resuspended in 200 µL of PBS. This volume was then used to intravenously inject approximately 3 million monocytes into the tail vein of a CD45.1 host mouse.

Soon after injection, a 1 mm wound was performed on the dorsal side of host mouse ear. At 14 days post wound induction, the wound skin was collected, and epidermis was separated by incubating with Dispase 5mg/mL at 37°C for 30 minutes. The epidermis was then immunostained for CD207 and CD45.2 following the same procedure described in the whole-mount staining section. Finally, the stained tissue was imaged using a STELLARIS 5 Leica confocal laser microscope.

### Flow cytometry

The epidermis of day 13 wounded CCR2-GFP mice was collected and separated from the dermis by incubation with 5mg/mL at 37°C for 30 minutes. The epidermis was then minced to a single cell suspension using fine point scissors inside FACS buffer (1x PBS, 3% FBS, 2mM EDTA, 0.05% Sodium Azide). The cell suspension was then filtered through a 70 µm and a 45 µm strainer, consecutively. Cells were washed with PBS and distributed in 1.5 mL Eppendorf tubes for staining. All staining steps were performed in a 100 µL volume in the dark on ice. Samples were first incubated with Zombie NIR Fixable Viability Kit (1:400, Biolegend; 423105). Cells were washed once with FACS buffer followed by incubation with Fc block reagent (BDBioscience; 553142) for 15 minutes. The following antibodies were mixed and added directly to the cell suspension: BUV395 CD45 (1:150, eBioscience; 363-0451-82), BV510 CD11b (1:150, Biolegend; 101235), BV421 Ly6C (1:150, Biolegend; 128031).

Cells were incubated for 30 minutes and washed using FACS buffer. Cells were fixed and permeabilized following the manufacturer’s instructions (BD Cytofix/Cytoperm kit, BDB554714). Cells were then resuspended in BD Perm/Wash buffer with AF647 CD207 (1:150, Biolegend; 110720) for 30 minutes. Cells were washed twice with BD Perm/Wash buffer followed by resuspension in a final volume of 250 µL FACS buffer for flow cytometry analysis.

Bone marrow MACS enriched monocytes were washed in PBS and resuspended in LIVE/DEAD Fixable Blue Dead Cell Stain Kit (1:500, ThermoFisher, L23105) for 30 min. Cells were washed once with FACS buffer followed by incubation with Fc block reagent (BDBioscience; 553142) for 15 minutes. The following antibodies were mixed and added directly to the cell suspension: PE CD3ε (1:150, BioLegend; 100307), PE CD45R (1:150, BioLegend; 103207), PE CD49b (1:150, BioLegend; 108907), PE Ly6G (1:150, BioLegend; 127607), PE NK1.1 (1:150, BioLegend, 108707), APC SiglecF (1:150, Biolegend; 155507), and FITC Ly6C (1:150, Biolegend; 128005). Cells were incubated for 30 minutes and washed using FACS buffer. Cells were resuspended in a final volume of 250 µL FACS buffer for flow cytometry analysis.

All samples were collected and unmixed using a Cytek Aurora spectral flow cytometer (Cytek Biosciences), with the Cytek SpectroFlo software (version 4.0.3). Flow cytometry data were analyzed using FCSExpress software (version 7.30.0030).

### ELISA

Homeostasis and wound tissue were collected using a punch biopsy and total protein was extracted using RIPA buffer (Thermo Scientific; 89901) and homogenized using fine point scissors. Extracted total protein was quantified using BCA Protein assay (Thermo Scientific; A55860). Samples were normalized to 1mg/mL of total protein using the corresponding ELISA kit assay diluent buffer. Day 2 extracted protein samples were run in duplicate using mouse ELISA kits for CXCL1 (Sigma; RAB0117), CXCL2 (Sigma; RAB0128), CXCL3 (Abcam; ab272191), CXCL5 (Sigma; RAB0131), and CXCL7 (Sigma; RAB0136). Day 7 wound tissue was collected as full thickness, epidermis only, or dermis only. Day 7 extracted protein samples were run in duplicate using mouse ELISA kits for CSF1 (Sigma; RAB0099), and IL-34 (ELK biotechnology; ELK5736). Statistical analysis and curve fitting were performed using GraphPad Prism (version 10.0). Raw optical density (OD) values obtained from the microplate reader were first corrected by subtracting the mean OD of the blank control wells. To generate the standard curve, a four-parameter logistic (4PL) non-linear regression model was applied to the log-transformed concentrations of the standard samples. Unknown sample concentrations were back calculated from the linear range of the standard curve using the interpolated value function in GraphPad Prism. Samples with OD values falling outside the lower limits of the standard curve’s linear range were marked as zero. Final concentration values were multiplied by the respective dilution factors to determine absolute protein concentration.

### Statistics and reproducibility

Data are expressed either as absolute numbers or percentages ± s.d. An unpaired, two-tailed Student’s *t*-test was used to analyze datasets with two groups. A paired, two-way ANOVA was used to analyze datasets across multiple time points. Actual P-values were provided in the legends when available. NS denotes no significance. Statistical calculations were performed using Prism v.10.4.0 (GraphPad). No statistical method was used to pre-determine sample size (n). Sample sizes are represented as distinct biological replicates in all experiments. Panels showing representative images are representative of at least two to five independent experiments, as indicated in the figure legends. \**P*<0.05, \*\**P*<0.01, \*\*\**P*<0.001 and \*\*\*\**P*<0.0001 indicate significant differences.

## Data and code availability

Bulk RNA-seq and single-cell RNA-seq data generated in this study have been deposited in the Gene Expression Omnibus under accession numbers GSE338807 and GSE338808, respectively. All other data that support the findings of this study are available from the corresponding author upon request.

## Authors contributions

A.D.S.-S. and S.P. designed the experiments and wrote the manuscript. A.D.S.-S. performed two-photon imaging, IMARIS analysis, chemokine receptor inhibition, whole-mount staining, scRNA-seq, flow cytometry, adoptive transfer, and ELISA assays. N.B. performed adoptive transfer, ELISA assays, and whole-mount staining. S.M. and H.K. performed flow cytometry. N.B. and A.B. performed image analysis and qRT–PCR. A.D.S.-S., N.B., S.M., and A.B. maintained mouse lines for all experiments. C.M.-M. performed bulk RNA-seq. M.-S.S. and G.W.L. provided critical feedback on the research and manuscript.

## Acknowledgments

We thank V. Greco for providing the Lang-EGFP, K14-H2B-mCherry, and K14-H2B-Cerulean mouse lines. We are grateful to L. Terrian and D. Bowman for their discussion and guidance on scRNA-seq analysis. We thank the MSU Genomics Core for library preparation and the University of California, Berkeley, QB3 Genomics Core for library sequencing. The data presented herein were obtained using instrumentation in the MSU Flow Cytometry Core Facility (RRID:SCR_026903). We also thank the MSU IQ Microscopy Core for instrument access and maintenance, and the MSU Campus Animal Resources for animal husbandry support. A.D.S.-S. was supported in part by the MSU John A. Penner Endowed Research Fellowship. This work was supported by the National Institutes of Health (NIH) under award numbers R01AR083086 and R01AI134696, by the MSU Discretionary Funding Initiative, and by the Department of Medicine Seed Funding.

---

**Figure S1. Epithelial cell migration on Day 2 post-injury. A**, Experimental design for the longitudinal study of in-vivo wound healing. Revisit imaging (camera icon) was performed on Day 0 and 5. Timelapse imaging (video icon) was performed on Day 2. **B**, Schematic of wound healing multiphoton intravital microscopy. **C**, Imaris track analysis of epithelial cells (Figure 1B) 2 days after wound induction. Colors project time (blue, 0h; red, 6h). *Top*: *x-y* view. *Bottom*: *x-z* view. Representative image from 3 mice. Scale bars, 100 µm. **D**, Zoomed migration tracks from **C**. The green frame is from the wound leading-edge epithelial migration zone, and the teal frame is from the epithelial proliferation zone^3^. Representative image from 3 mice. Scale bars, 100 µm. **E**, Mean total displacement of individual epithelial cells tracks over 6h plotted as a function of distance from the wound. Dashed line, initial wound boundary. Data analyzed using paired two-way ANOVA; *n* = 3 mice; data are mean ± s.d. with each dot representing individual mice. *** *P* < 0.001, **** *P* < 0.0001.

**Figure S2. Tamoxifen-induced CreER recombination in vivo. A**, The schematic illustrates the experimental design for CreER-mediated recombination, including the timing of tamoxifen injection and subsequent in vivo imaging in mice, such as in Epi-*Rac1* (*K14-CreER;Rac1^−/−or^ ^+/−^*) and LC-*Rac1* (*huLangerin-CreER Rac1^−/−or^ ^+/+^*), dual-labeling mouse line (*huLangerin-CreER; Rosa-stop-tdTomato; Lang-EGFP; K14-H2B-mCherry*). Revisit imaging (camera icon) was performed on Day 0 and 5. Timelapse imaging (video icon) was performed on Day 2. **B**, *In-vivo* microscopy images show *x-y* view of epithelial cells (red nuclei) in the epidermis of control and Epi-*Rac1* KO mice at 0, 5 and 8 days after wound induction. Dashed line indicates initial wound boundary or scab. Representative image from 3 mice per group. Scale bars, 100 µm. **C**, Stereomicroscopy imaging of control and Epi-*Rac1* KO mice at Day 0 and 8 wound healing. Representative image from 3 mice per group. Scale bars, 100 µm.

**Figure S3. The majority of MHC-II⁺ cells in the wound epidermis are Langerhans cells. A**, Confocal immunofluorescent images of MHC-II+ Langerhans cells at the wound epidermis at days 7, 13 and week 3. *Left:* Images show *x-y* view of MHC-II+ cells (green), Langerhans cells (red, CD207), and cell nuclei (blue, DAPI). *Right*: green and red composite and zoomed view. Dashed line indicates initial wound boundary. Representative images from 3 mice. Scale bars, 100 µm. **B**, Percentage of Langerhans cells (CD207+) positive for MHC-II+ in the epidermis during homeostasis and days 7, 13, and week 3 of wound healing. Data analyzed using unpaired one-way ANOVA; *n =* 3 mice; data are mean ± s.d. with each dot representing individual mice.

**Figure S4. IL-34 is increased on Day 7 of wound healing. A**, ELISA assay of CSF1 present at the wound site on Day 7 of healing compared to homeostasis. **B**, ELISA assay of IL-34 present at the wound site on Day 7 of healing compared to homeostasis. **C**, ELISA assay of CSF1 present at Day 7 of wound healing in the epidermis compared and dermis. **D**, ELISA assay of IL-34 present at Day 7 of wound healing in the epidermis compared and dermis. **A-D**, Data analyzed using unpaired two-tailed *t*-test; *n = 4* mice; data are mean ± s.d. with each dot representing individual mice. * *P* < 0.05.

**Figure S5. Langerhans cells quickly recover their normal distribution in the epidermal basal layer after wound induction. A**, *In-vivo* microscopy images of epithelial cells (red nuclei) and LCs (green) in the epidermis 10 days after wound induction. The wound and far zones are as designated in (Figure 3A). The dashed line separates the basal and granular layers of the epidermis. The solid line separates the epidermis and dermis. *Left*: *xz* and *xy* views of the wound epidermis. *Right*: *xy* and *xz* views of the far epidermis. Representative images from 6 mice. Scale bars, 50 µm. **B-C**, Timeline of LC vertical distribution in the epidermis during wound healing at the wound (**B**) and far (**C**) zones. Data quantified as the minimal distance to the basement membrane (solid line). *n* = 6 mice. **D-E**, Timeline of LC horizontal distribution in the epidermis during wound healing at the wound (**D**) and far (**E**) zones. Data quantified as the average minimum distance to 3 closest LCs. *n* = 6 mice. **B**-**E**, Representative quantification of LC distribution. Data analyzed using paired one-way ANOVA; data are mean ± s.d. with data points representing individual cells. **** *P* < 0.0001.

**Figure S6. scRNA-Seq heterogeneity in the epidermis on Day 11 after wound induction. A**, UMAP of scRNA-Seq data enriched for LCs collected from the ear epidermis 11 days after wound induction. LC clusters were identified by high *Langerin* (*Cd207*) expression and were divided into activated (high Cd80), steady-state (low Cd80), cycling (Top2a), migrating (Ccr7), and apoptotic (Ddias) categories. Other clusters were identified according to their top DEGs. **B**, Violin plots of scRNA-Seq data after light quality control showing each cluster’s total read counts, feature gene counts, and percentage of mitochondrial genes expressed. **C**, *Left*: Violin plots showing the expression of cluster-defining genes across each cluster. *Right*: Feature plots showing the expression of the gene mentioned in the same row. Mono, Monocytes. Act, activated. Ss, steady-state. Cyc, cycling. Mig, migrating. Apop, apoptotic. KC, keratinocytes (epithelial cells). Neut, neutrophils. T-mem, T memory cells.

**Figure S7. eLC subclustering and gene expression of genes related to GO terms. A**, UMAP of monocyte and Langerhans cells re-clustering, highlighting eLC subclusters. eLC were subclustered into steady-state (low *Cd80*, high *Cdh1*) and activated (high *Cd80*, low *Cdh1*). **B**, *Left*: violin plots showing *Cd80* and *Cdh1* expression in eLC subclusters. *Right*: Feature plots showing the expression of the gene mentioned in the same row. Data analyzed using an unpaired *t*-test. **** *P* < 0.0001. **C-F**, Dot plots showing the expression of genes in (Fig. 5g) GO terms according to clusters: Mono (**C**), pre-mLC (**D**), mLC (**E**), and eLC (**F**). Highlighted color shades match genes to their corresponding GO term.

**Figure S8. Adoptive transfer of monocytes give rise to Langerhans cells during wound healing. A**, The schematic shows the experimental design for adoptive transfer of monocytes to test their differentiation into Langerhans cells during wound healing. **B**, Flow cytometry gating strategy for the enrichment of bone marrow harvested monocytes. **C**, Mean percentage of monocytes harvested before and after MACS isolation. Data analyzed using unpaired two-tailed *t*-test; *n =*3 mice; data are mean ± s.d. with each dot representing individual mice. **** *P* < 0.0001. **D**, Confocal immunofluorescent images of donor monocytes-derived Langerhans cells at the wound epidermis 14 days after wound induction. *Left*: Images show *x-y* view of LCs (green, CD207), donor origin (red, CD45.2), and cell nuclei (blue, DAPI). *Right*: zoomed view in composite, green channel only, and red channel only. Dashed line indicates initial wound boundary. Representative images from 3 mice. Scale bars, 100 µm.

**Figure S9. Characterization of epidermal CCR2-GFP cells at Day 13 of wound healing. A**, Flow cytometry gating strategy for the characterization of CCR2-GFP myeloid cells in the epidermis at Day 11 of wound healing. **B**, Mean percentages of epidermal LyC6+, Ly6C+CD207+, and CD207+ cells in CCR2-GFP negative and positive myeloid cells at Day 11 of wound healing. Data analyzed using unpaired two-way ANOVA; *n* = 4 mice; data are mean ± s.d. with each dot representing individual mice. * *P* < 0.05.

**Figure S10. CXCR4 and CCR7 are not required for Langerhans cell migration during re-epithelialization. A**, *In-vivo* microscopy images show *x-y* view of dermis SHG collagen (gray) and LCs (green) in the epidermis 5 days after wound induction. Dashed line indicates initial wound boundary. *Top left*: control mouse. *Top right*: *LC-Cxcr4 KO* mouse. *Bottom*: *Ccr7 KO* mouse. Representative images are shown. n = 3 control, n = 3 LC-Cxcr4 KO, and n = 4 Ccr7 KO mice. Scale bar, 50 µm. **B**, Mean LC number inside the wound epidermis from control, LC-*Cxcr4 KO*, and *Ccr7 KO* mice. Imaging was performed 5 days after wound induction. Wound area quantified 0.09 mm^2^ per mouse. Data analyzed using unpaired one-way ANOVA; n = 3 control, n = 3 LC-Cxcr4 KO, and n = 4 Ccr7 KO mice; data are mean ± s.d. with each dot representing individual mice.

## Supplementary Video Legends

**Video S1. Day 2 epithelial and LC migration.** Time-lapse recording over 6h of epithelial cells (red nuclei) and LCs (green) 2 days post-wound induction (PWI). Dashed line, initial wound boundary. Solid line, basal membrane separating epidermis from dermis. *Top*: *x-y* view. *Bottom*: *x-z* view shows the epidermis (red) and dermis (collagen SHG, blue). Representative video from 3 mice.

**Video S2. Epi-*Rac1* KO and control mice Day 2 epithelial and LC migration.** Time-lapse recording over 6h in *x-y* view of epithelial cells (red nuclei) and LCs (green) 2 days post-wound induction (PWI). *Top*: control mouse. *Bottom*: Epi-*Rac1 KO* mouse. Dashed line, initial wound boundary. Representative video from 3 mice.

**Video S3. In vivo LC proliferation within the wound epidermis at Day 7 post injury.** Time-lapse recording over 5h in *x-y* view of proliferative LCs (green) at 7 days post-wound induction (PWI). Dermis SHG collagen is shown in gray. White arrows indicate diving LCs. Dashed line, initial wound boundary. Representative video from 3 mice. Scale bar, 50 µm.

**Video S4. Zoom of LC proliferation on Day 7 post injury.** Zoomed-in example of proliferative LCs (green) in **Video S3**. Representative video from 3 mice. Scale bar, 10 µm.

**Video S5. Day 2 epithelial cell and LC migration in Cxcr2-inhibited and control mice.** Time-lapse recording over 6h in *x-y* view of epithelial cells (red nuclei) and LCs (green) 2 days post-wound induction (PWI). *Top*: control mouse 1% DMSO. *Bottom*: drug-treated mouse CXCR2 (Danirixin). Dashed line, initial wound boundary. Representative video from 3 mice.

