## Supplemental Figures for "Dual lineages of Langerhans cells cooperate to restore the immune barrier after skin injury"

Figure S1. Epithelial cell migration on day 2 after injury.

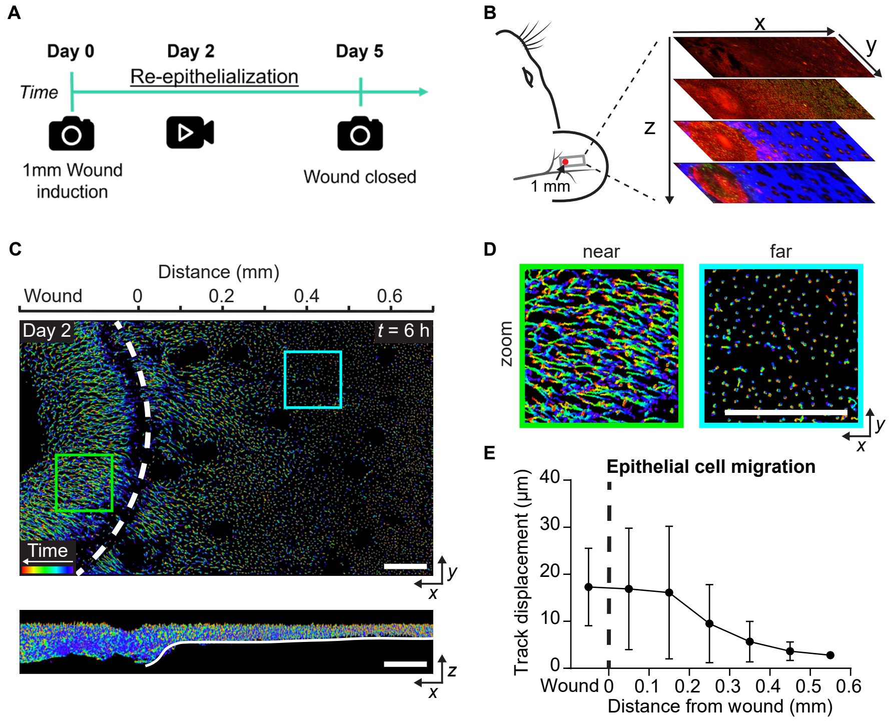

Figure S2. Tamoxifen-induced CreER recombination in vivo.

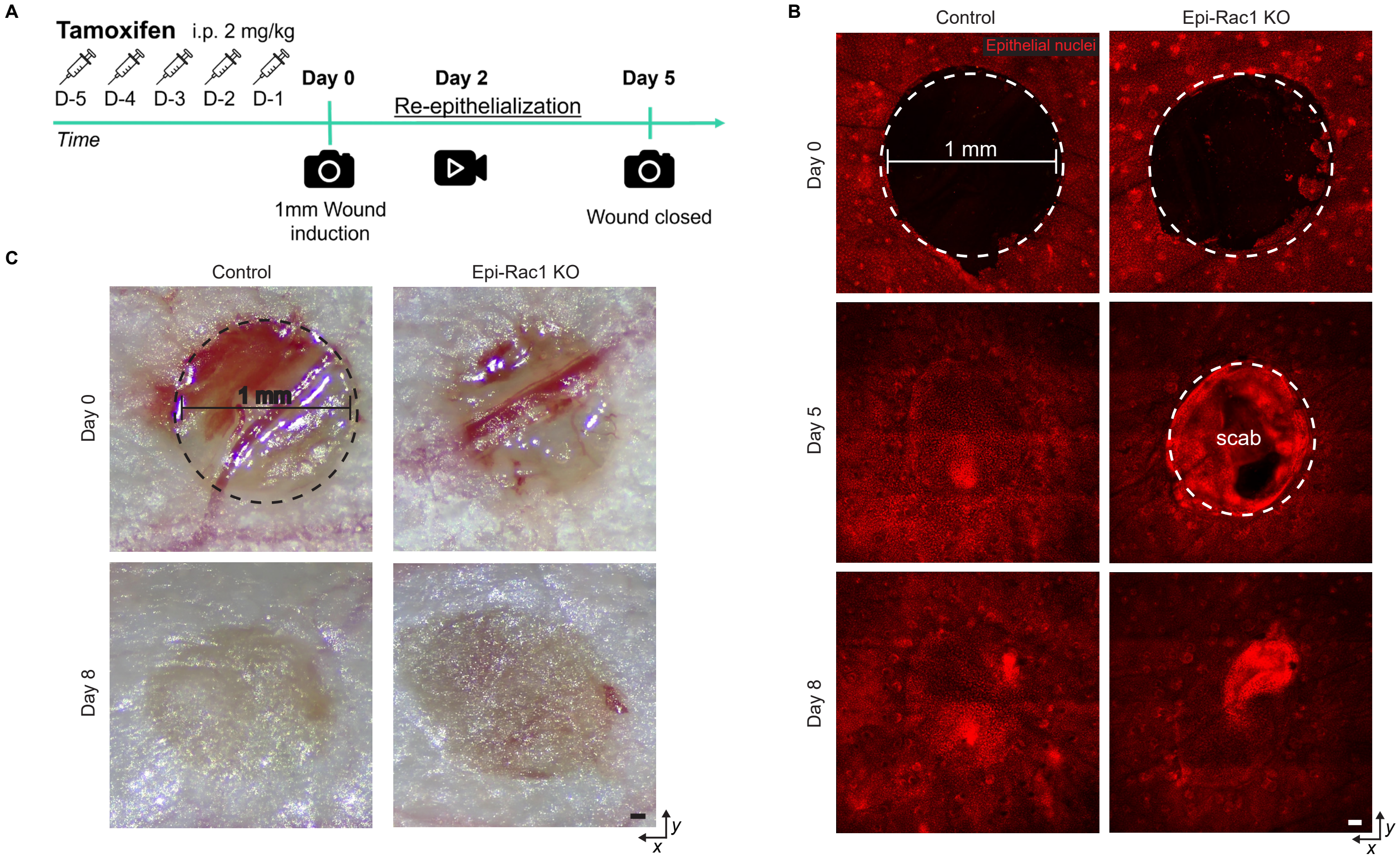

Figure S3. The majority of MHC-II$^{+}$ cells in the wound epidermis are Langerhans cells.

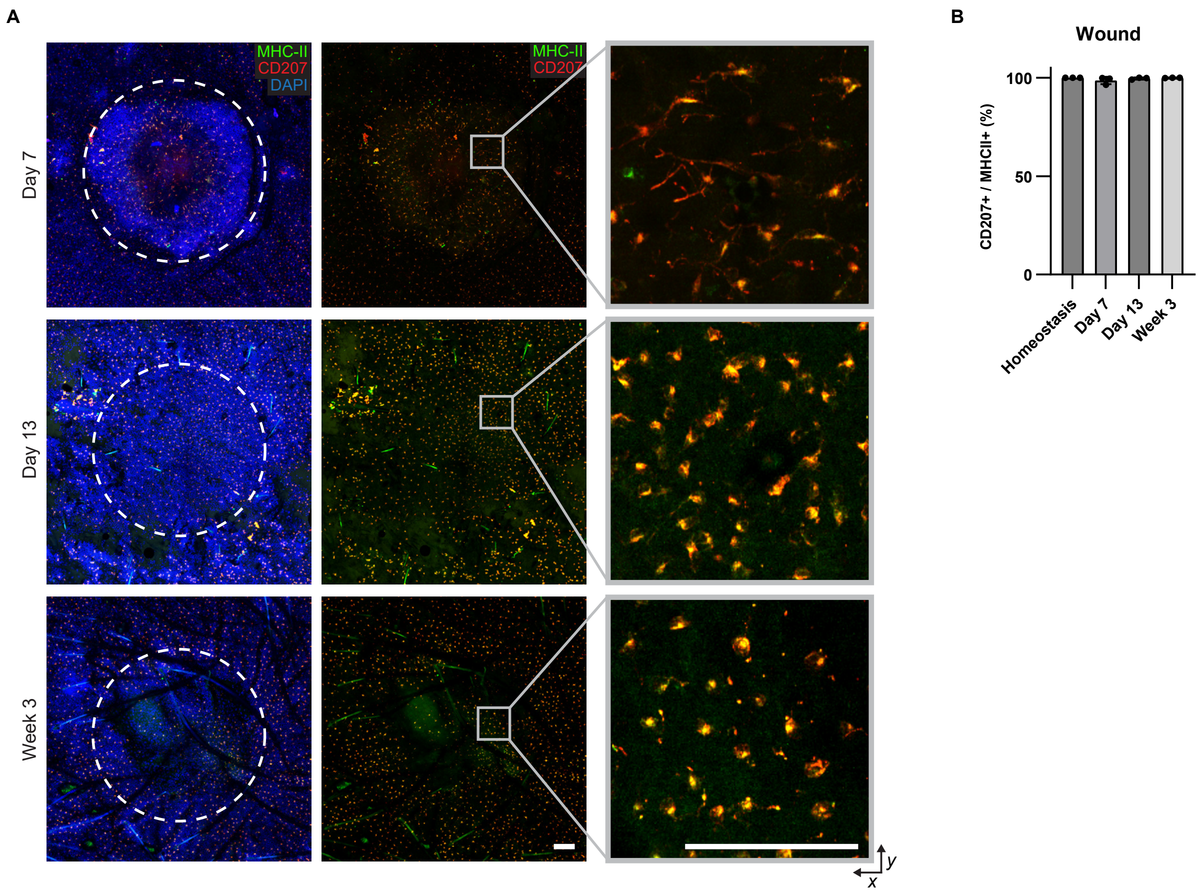

Figure S4. IL-34 is increased on day 7 of wound healing.

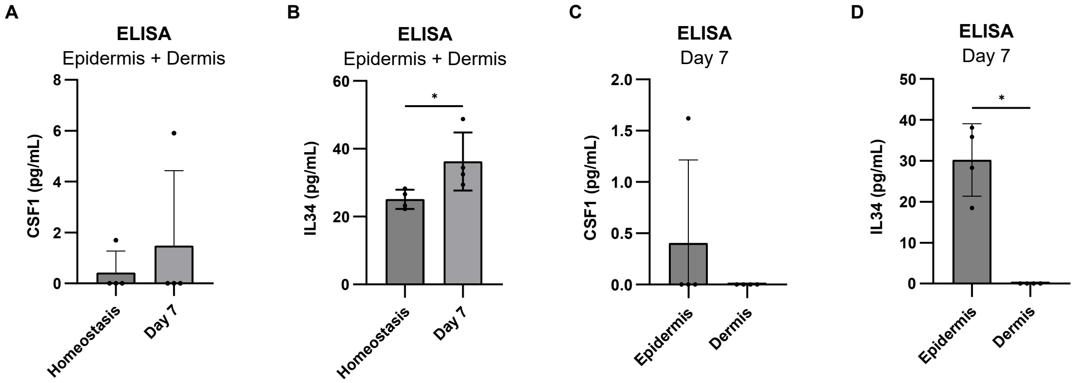

Figure S5. Langerhans cells quickly recover their normal distribution in the epidermal basal layer after wound induction.

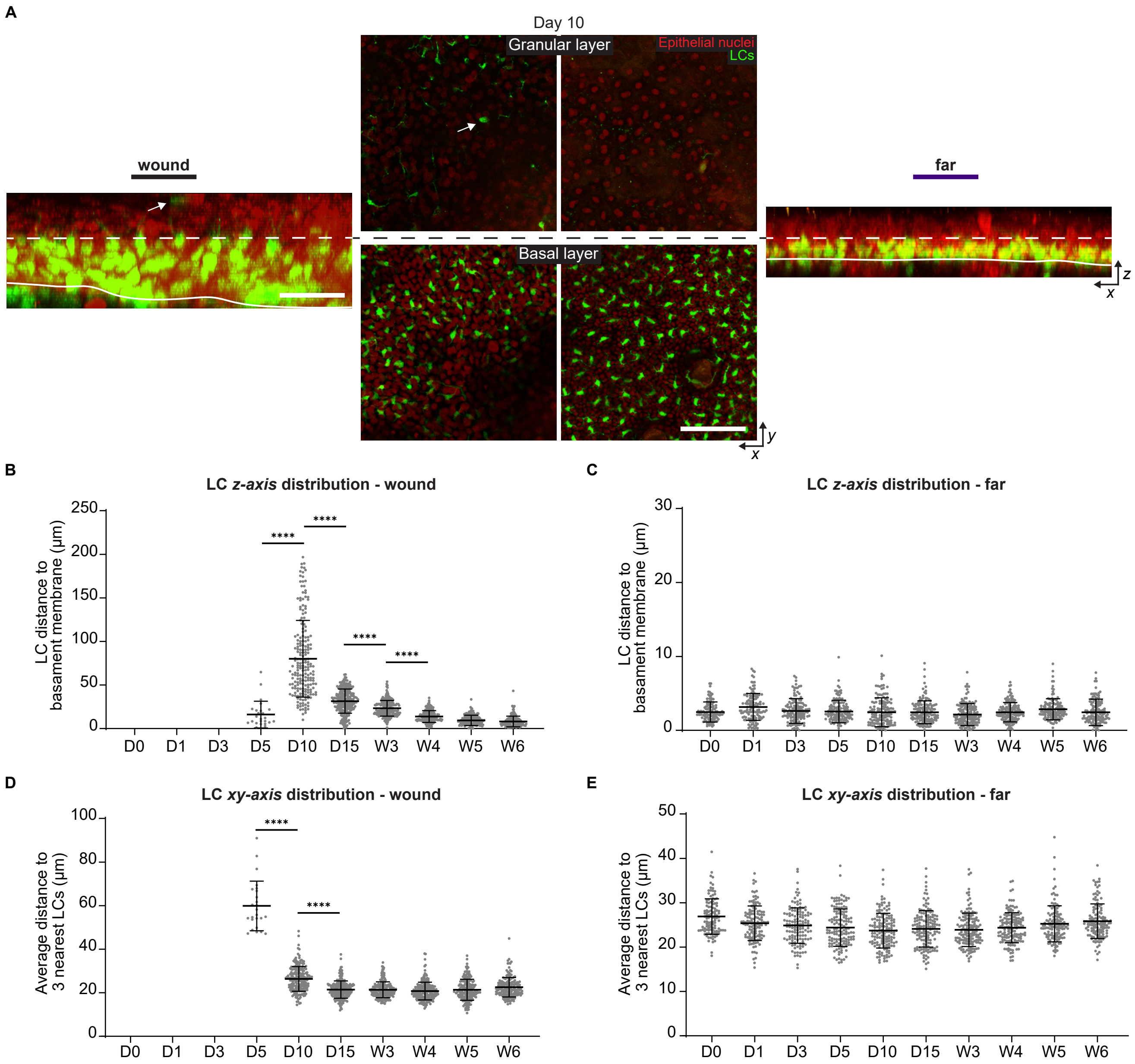

Figure S6. scRNA-Seq heterogeneity in the epidermis on day 11 after wound induction.

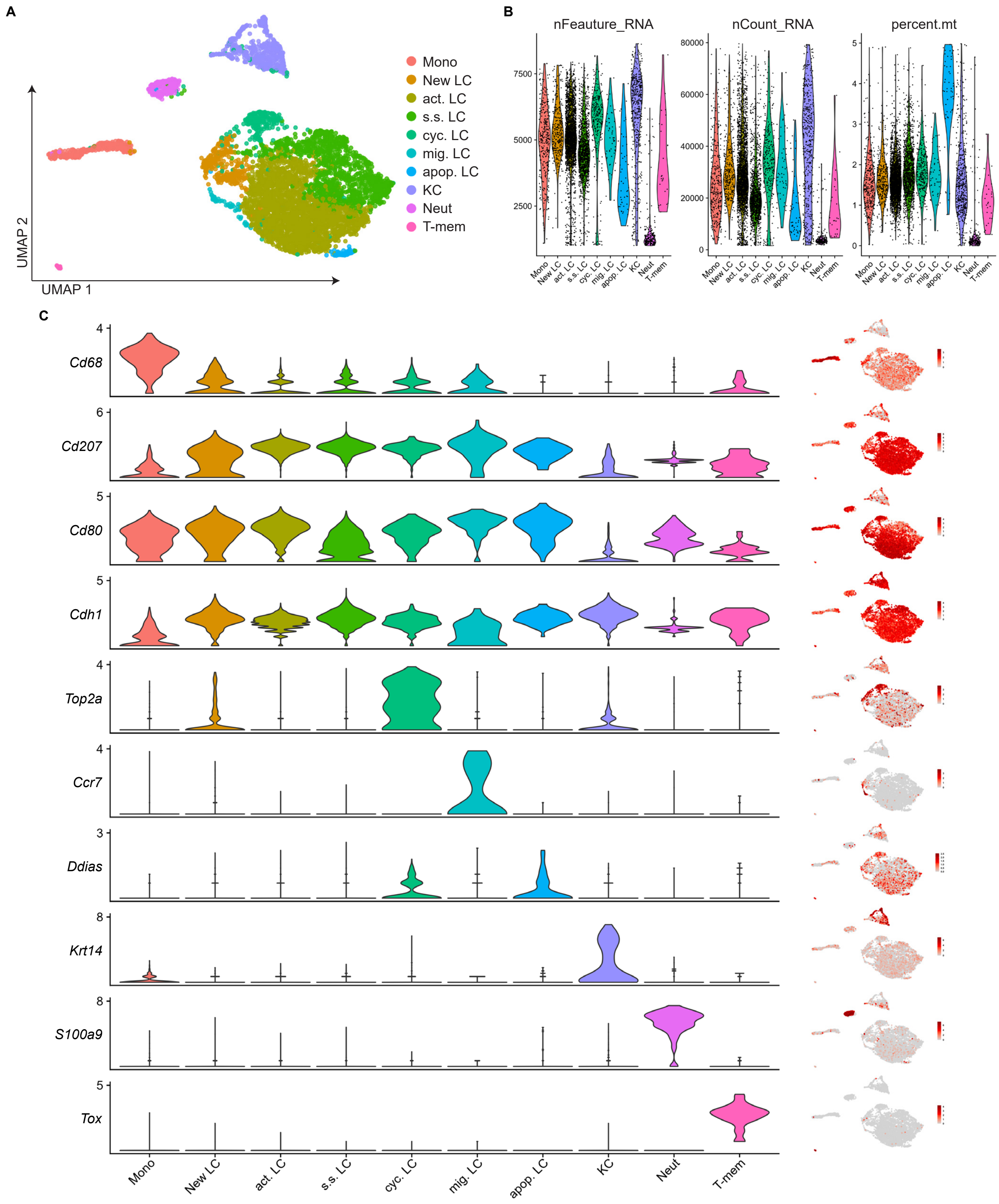

Figure S7. Steady-state vs activated eLC and GO terms gene expression.

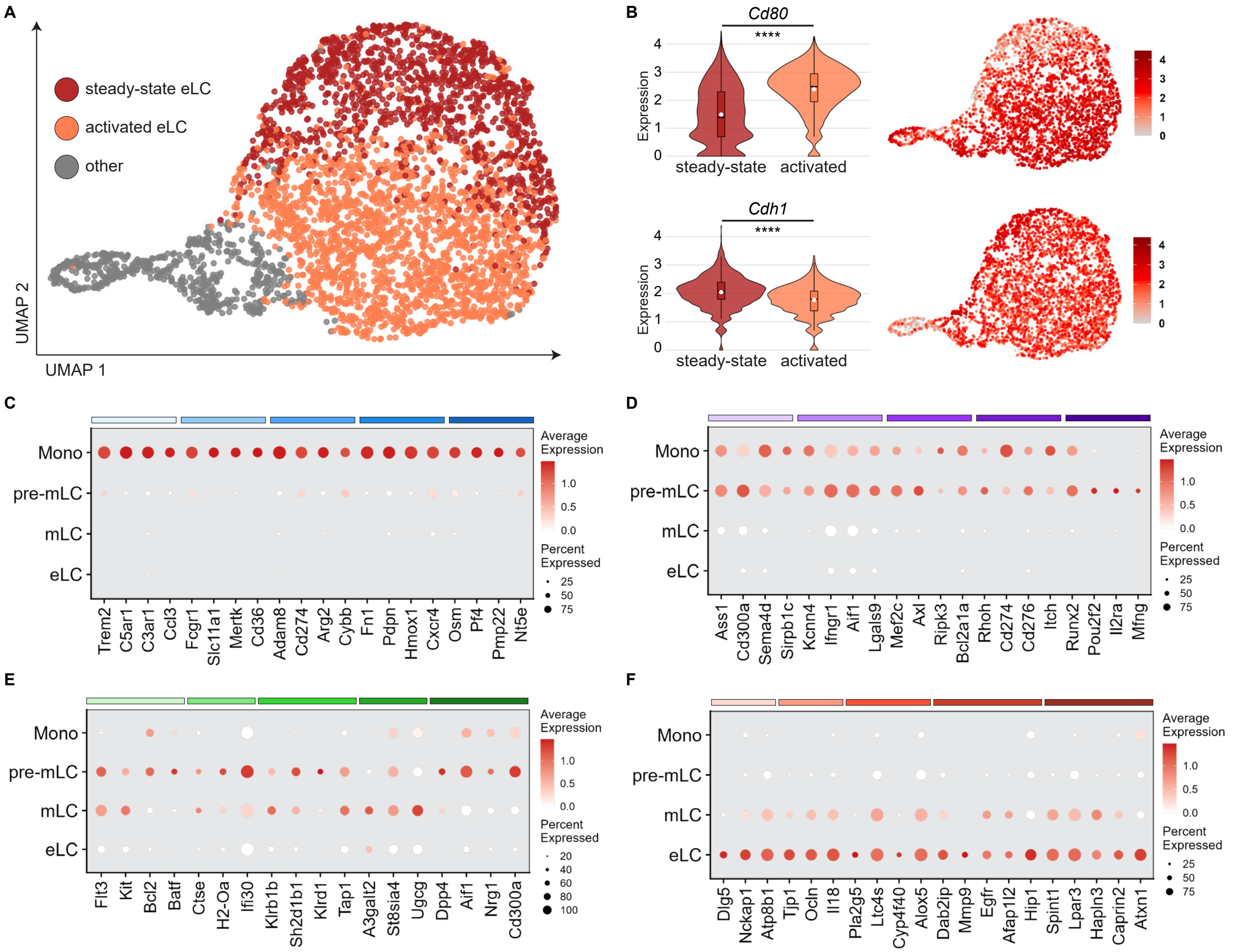

Figure S8. Adoptive transfer of monocytes give rise to Langerhans cells during wound healing.

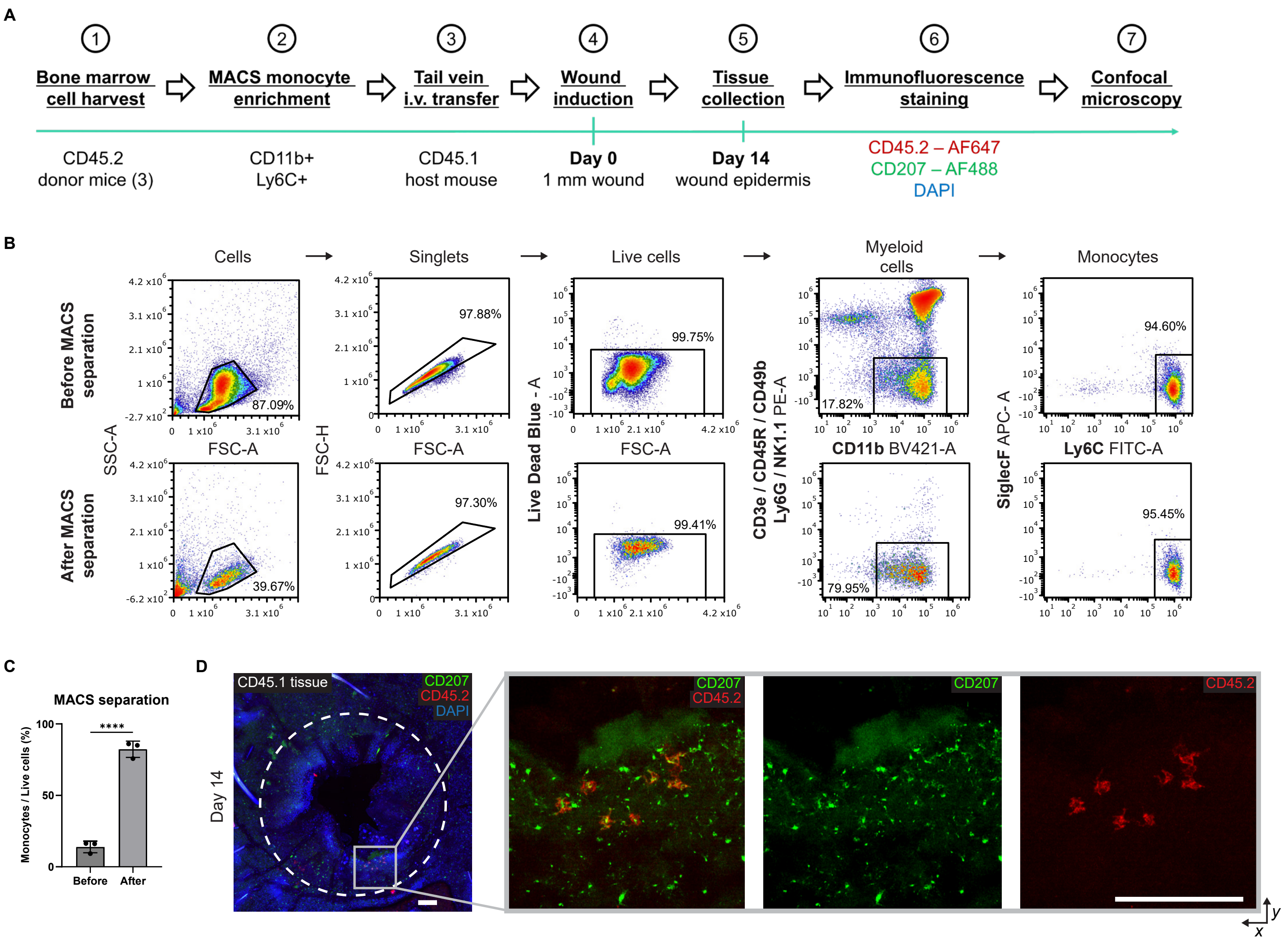

Figure S9. Characterization of epidermal CCR2-GFP cells at day 13 of wound healing.

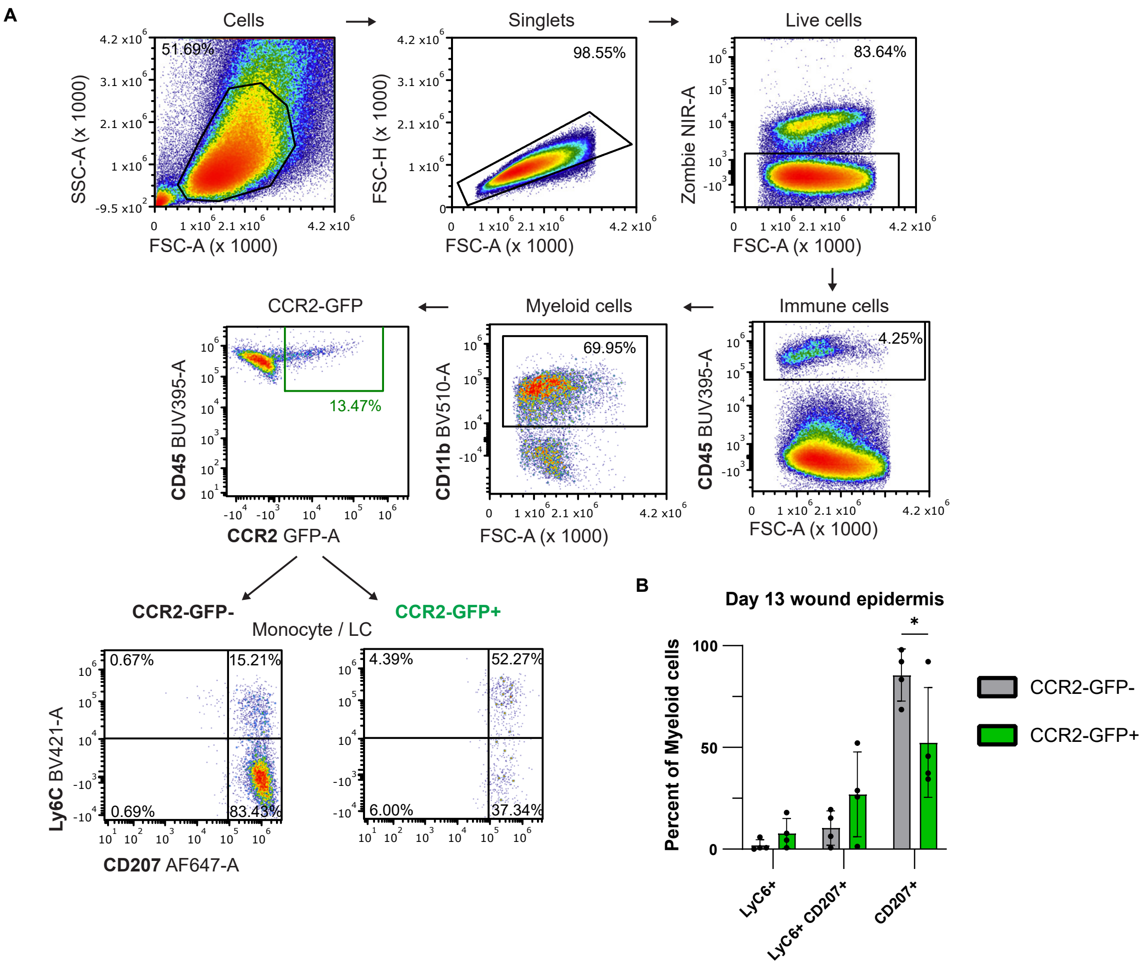

Figure S10. CXCR4 and CCR7 are not required for Langerhans cells migration during re-epithelialization.

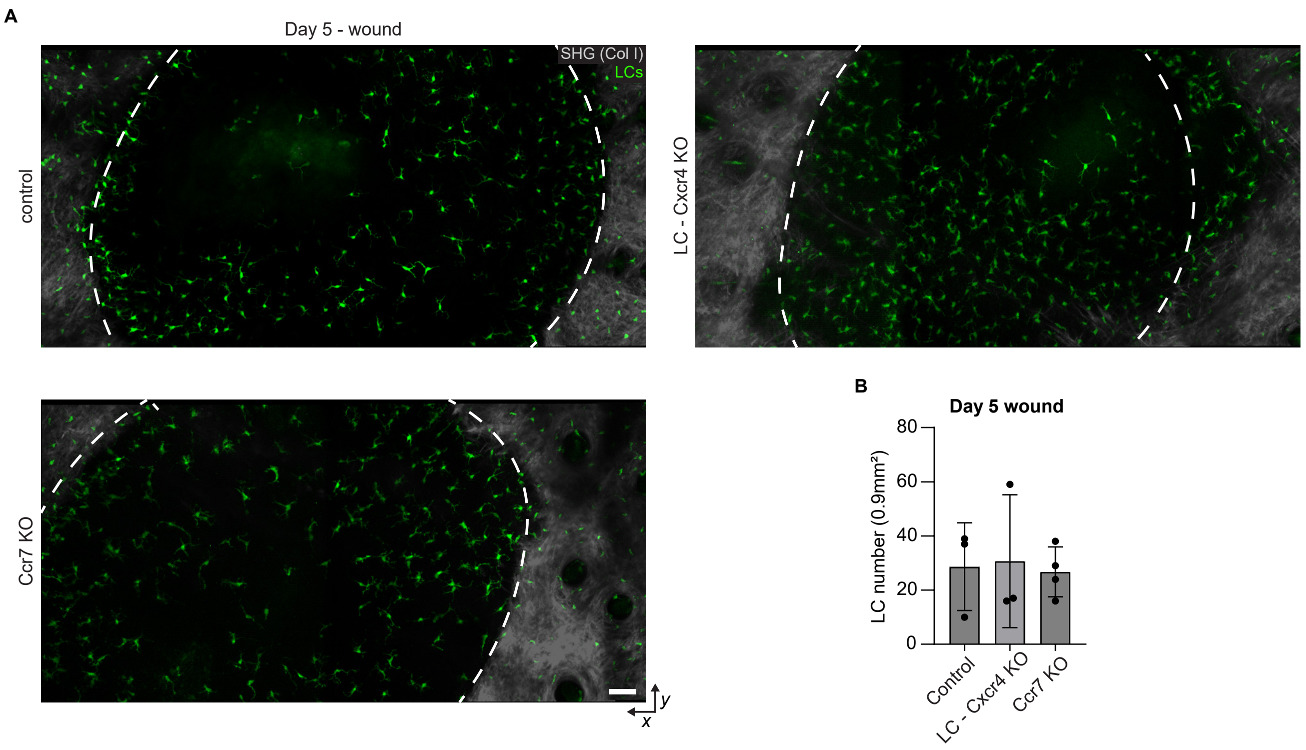
